# Vestigial Auriculomotor Activity Indicates the Direction of Auditory Attention in Humans

**DOI:** 10.1101/786525

**Authors:** Daniel J. Strauss, Farah I. Corona-Strauss, Andreas Schroeer, Philipp Flotho, Ronny Hannemann, Steven A. Hackley

## Abstract

It is commonly assumed that, unlike dogs and cats, we humans do not make ear movements when focusing our attention reflexively toward novel sounds or voluntarily toward those that are goal–relevant. In fact, it has been suggested that we do have a pinna–orienting system. Although this system became vestigial about 25 million years ago, it still exists as a “neural fossil” within the brain. Consistent with this hypothesis, we demonstrate for the first time that the direction of auditory attention is reflected in the sustained electrical activity of muscles within the vestigial auriculomotor system.

Surface electromyograms (EMGs) were taken from muscles that either move the pinna or alter its shape. To assess reflexive, stimulus-driven attention we presented novel sounds from speakers at four different lateral locations while the participants silently read a boring text in front of them. To test voluntary, goal-directed attention we instructed participants to listen to a short story coming from one of these speakers, while ignoring a competing story from the corresponding speaker on the opposite side.

In both experiments, EMG recordings showed larger activity at the ear on the side of the attended stimulus, but with slightly different patterns. Upward movement (perking) differed according to the lateral focus of attention only during voluntary orienting; rearward folding of the pinna’s upper-lateral edge exhibited such differences only during reflexive orienting. The existence of a pinna-orienting system in humans, one that is experimentally accessible, offers opportunities for basic as well as applied science. It could lead to a better understanding of the evolution of auditory attention and support the near real–time decoding of auditory attention in technical applications, for example, for attentionally controlled hearing aids that preferentially amplify sounds the user is attempting to listen to.

## Introduction

A review of research (Hackley, 2015) on pinna–orienting in humans identified three relevant findings scattered across the preceding 100–or–so years. The first was Wilson’s (1908) (Wilson, 1908) oculo-auricular phenomenon, in which shifting the gaze hard to one side elicits a 1 to 4 mm deflection of the lateral rim of both ears. The relevance to spatial attention is uncertain, though, because the relation between the side with the largest ear movement and that to which gaze is directed has been weak and inconsistent across studies (Gerstle & Wilkinson, 1929; Urban, Marczynski, & Hopf, 1993; O’Beirne & Patuzzi, 1999). Additional evidence comes from a 1987 study (Hackley, Woldorff, & Hillyard, 1987) of the bilateral postauricular muscle (PAM) reflex (onset latency = 10 ms) to acoustic onset transients. Increased amplitudes were observed when subjects directed their attention to a stream of tones on the same side as the recorded muscle while ignoring a competing, contralateral stream. Comparisons across left/right stimulus, attention, and PAM combinations localized modulation to the motor limb of the reflex arc. This pattern could indicate that the muscle behind an ear is primed when attention is directed toward that side. Finally, a 2002 experiment (Stekelenburg & Van Boxtel, 2002) found that the automatic capture of attention by unexpected sounds coming from a speaker hidden to the left of the participant elicited greater activity in the left than right PAM.

Apart from the research just described, functional studies of the human auriculomotor system have been mainly limited to the PAM reflex, in the context of audiometry or affective psychophysiology. The auriculomotor system lies essentially untouched in the literature. Here we present for the first time evidence that our brains retain vestigial circuitry for orienting the pinnae during both exogenous, stimulus-driven attention (to brief, novel sounds) and endogenous, goal-directed attention (to sustained speech). We also demonstrate a complex interplay of different auricular muscles which may be causally linked to subtle movements of the pinnae.

## Results

### Experiment 1 – Exogenous Attention

To examine automatic, stimulus-driven attention we used novel sounds similar to those of the just mentioned study (Stekelenburg & Van Boxtel, 2002) (e.g., traffic jam, baby crying, foot steps). However, we presented them randomly from four different speakers (at ±30°, ±120°; Fig. 1) rather than just one, while the subject read a boring essay. As we were interested in the interactive role of distinct muscles in attempting to shape and point the pinnae, we recorded EMG from posterior, anterior, superior, and transverse auricular muscles (PAM, AAM, SAM, and TAM). Visual evidence had previously been limited to still photos of Wilson’s oculo– auricular phenomenon (Wilson, 1908), so we supplemented our EMG data with videos from four high-definition cameras (see Methods and supplementary videos to Experiment 1). To confirm that our findings would generalize to the population most likely to benefit from attention-sensitive hearing aids, older (62.7 ± 5.9 y) as well as younger (24.1 ± 3.1 y) adults were tested.

**Figure 1:**
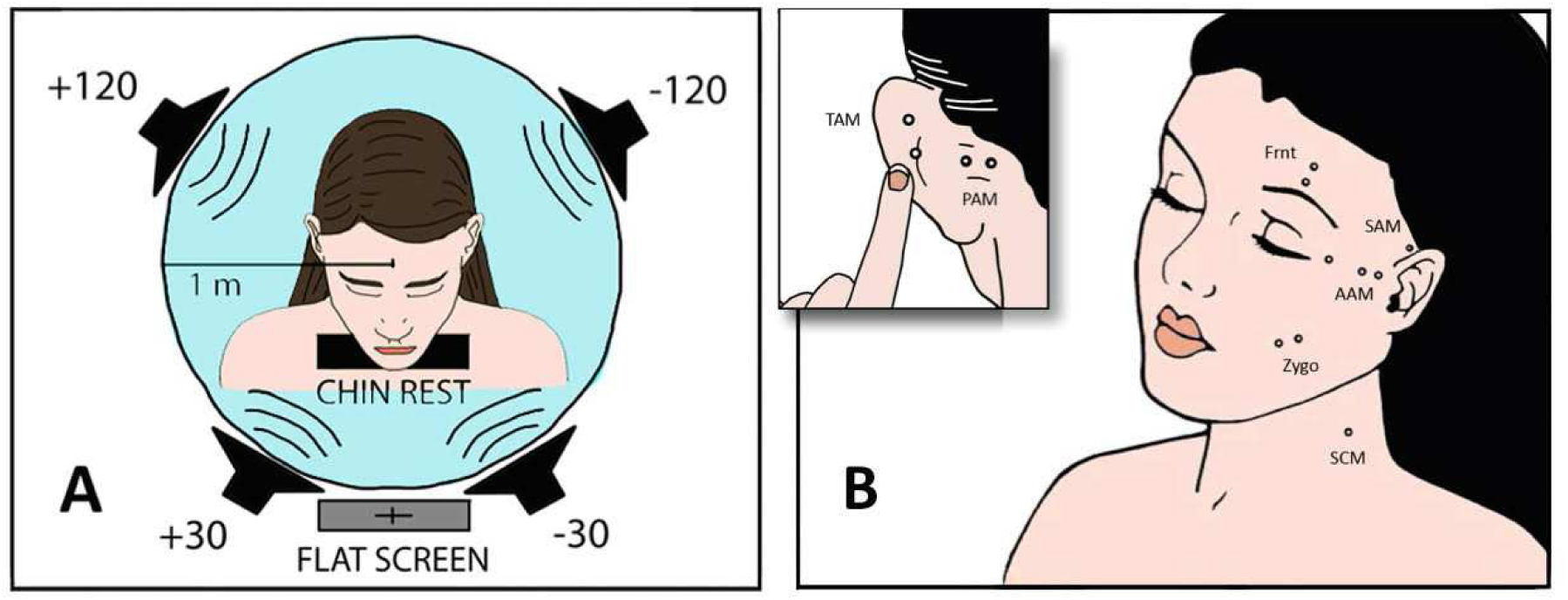
Experimental setup. (A) Four loudspeakers presented novel sounds (Exp. 1) or stories (Exp. 2) at 30° to the left or right of fixation or behind the interaural axis. Instructions, text, or fixation cross was displayed on a 55 in flat screen. (B) Surface EMGs were recorded bilaterally from four auricular muscles as well as from left zygomaticus major, frontalis, and sternocleidomastoideus, using a bandpass of 10 – 1000 Hz and a sampling rate of 9600 Hz. Separation of paired auricular electrodes was 1 cm.

The signal-averaged EMG waveforms of Fig. 2 show well–defined responses with an onset latency of about 70 ms, responses that vary in amplitude, duration, and morphology according to the relative direction of the sound source (The corresponding individual epoch data for each participant and stimulus can be found in Experiment 1 - figure supplement 2 for the PAM). Mean amplitudes were subjected to a mixed, repeated-measures analysis of variance, with factors of age group, stimulus-muscle correspondence (ipsi-/contralateral), and anterior/posterior stimulus direction. EMG amplitudes were larger for stimulus sources on the same side as the recorded ear for PAM, AAM, and TAM [*F* (1, 26) = 47.44, 17.01, and 47.53, respectively; *p*s < 0.001; 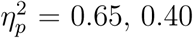, and 0.65] but not SAM.

**Figure 2:**
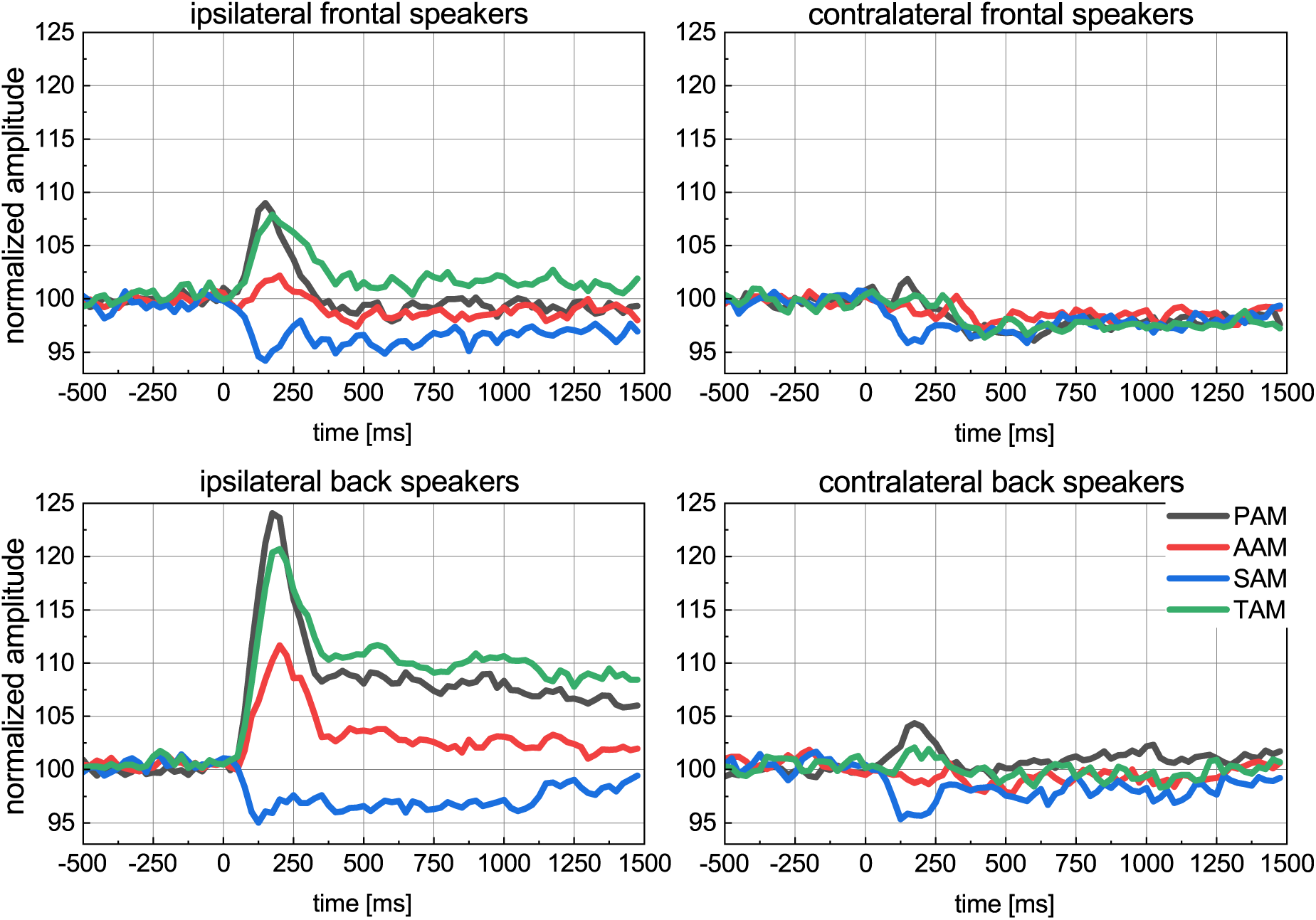
Experiment 1. Grand average (N = 28) of the baseline corrected and normalized eventrelated electromyograms at the four auricular muscles for the recording ipsilateral (left panel) and contralateral (right panel) to the stimulation; top: front speakers (30°), bottom: back speakers (120°). The contralateral-ipsilateral organization of our data set is justified by a preliminary analysis that obtained null effects for left–versus–right using a more complete factorial structure (left/right stimulus direction × left/right recording site).

Responses were also larger to sounds emanating from the back than the front speakers for PAM, AAM, and TAM [*F* (1, 26) = 32.1, 12.0, and 19.9, respectively; *p*s < 0.003, 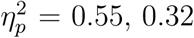, and 0.43]. Posterior, ipsilateral stimulation elicited the most vigorous responses from these three muscles. In particular, the interaction between the factors ipsi/contra and anterior/posterior for PAM, AAM, and TAM yields *F* (1, 26) = 40.4, 14.9, and 23.6, respectively; *p*s < 0.002; 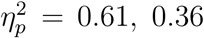, and 0.48. There were no interactions involving age group, but a main effect indicated that older participants had smaller AAM responses [*F* (1, 26) = 6.0, *p* < 0.03, 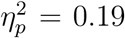]. These results support our hypothesis that the human brain retains circuits that attempt, futilely, to point the ears in the direction of unexpected, potentially relevant sounds. The corresponding vestigial auriculomotor drive appears to be causally linked to very small ear displacements, see Experiment 1 - figure supplement 1 and `Experiment - video supplement 1 and 2`.

We note, however, that increased AAM activity during the capture of attention by ipsilateral, *posterior* sounds was unexpected (Fig. 2). Perhaps in our remote ancestors PAM–AAM coactivation served to stabilize the base of the pinna and reduce occlusion of the ear canal during changes in ear shape and orientation. The suppression of SAM activity was also unexpected. However, a study of pinna orienting in cats, whose tall ears resemble those of prosimians, showed that they tend to tilt the top of the ear downward slightly when orienting to a lateral target (Fig. 3 in ref. (Populin & Yin, 1998)).

**Figure 3:**
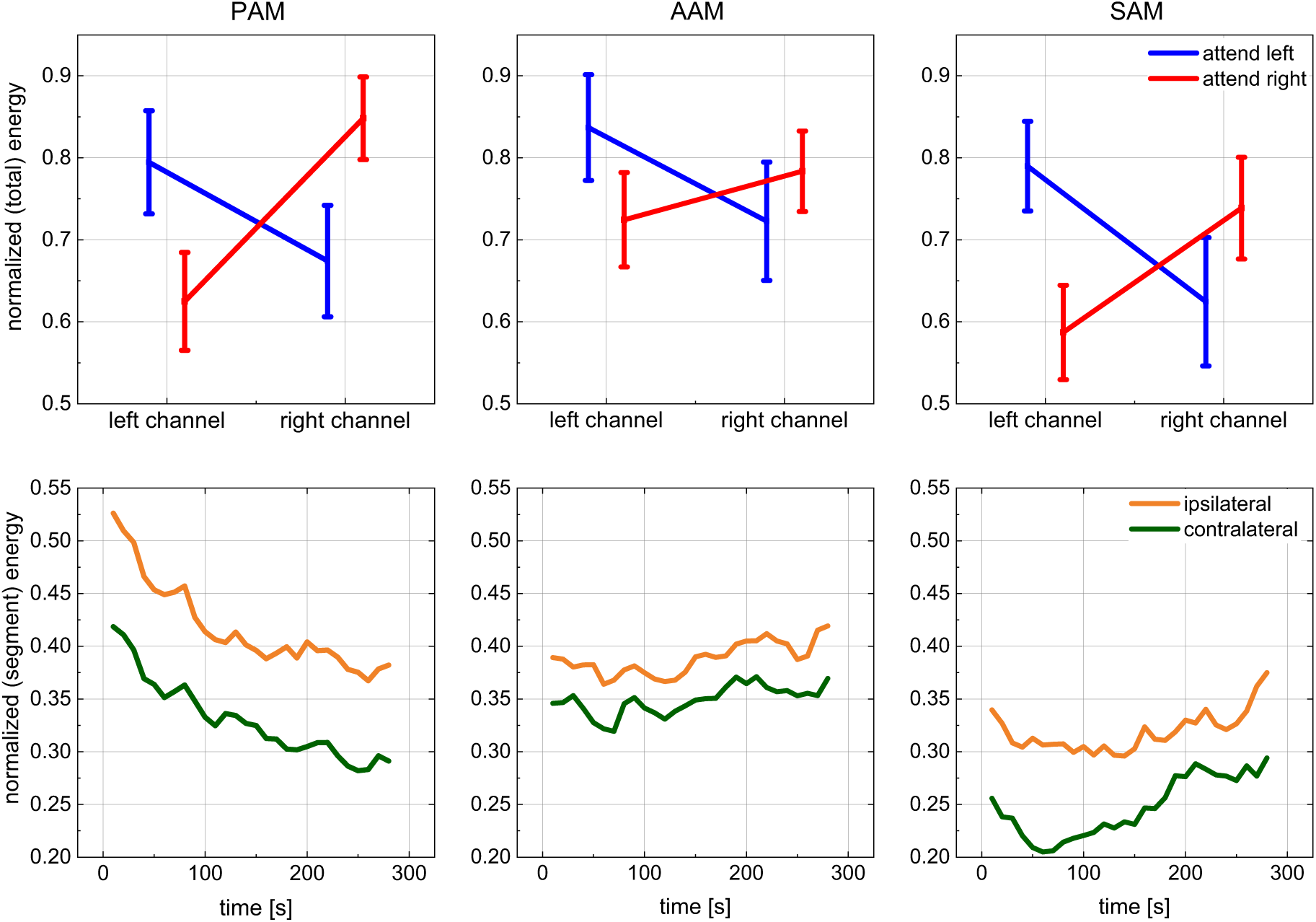
Experiment 2. Grand average of the PAM, AAM, and SAM activity when stories were played from the back speakers (±120°). Top: normalized (total) energy of the left/right recording channels during attention to the left or right story (bars represent the standard error). Bottom: time– resolved activity (each sampling point represents the energy induced in consecutive 10 s segments) after pooling the ipsi– and contralateral signals with a segment–wise normalization. For corresponding data for the front speakers, see Experiment 2 - figure supplement 1.

Two alternative interpretations of the ipsi– versus contralateral main effect should be considered. The first is that participants may have oriented their gaze toward the unexpected sounds, thereby triggering Wilson’s (1908) oculo–auricular phenomenon. To test this hypothesis, we signal averaged epochs of horizontal EOG using the same analysis parameters as described above for auricular EMG. Voltages were converted to degrees of arc separately for each participant, based on findings from a cursor-tracking protocol (±35°). Grand average waveforms (Experiment 1 - figure supplement 3-6) show a complete absence of systematic horizontal movements in the relevant time period. Furthermore, the 70-ms onset latencies we observed for auricular EMG responses is beyond the range of all but the very fastest saccades (Fischer & Ramsperger, 1984).

A second alternative interpretation is that participants oriented not with their ears or eyes, but by lifting their chin off the chin rest and rotating their head toward the novel sound. If humans have a vestibulo-auricular reflex as do cats (Tollin, Ruhland, & Yin, 2009), such head rotations could have indirectly triggered activity in the auricular musculature. This interpretation can be dismissed for three reasons: (1) Video monitoring indicated excellent compliance with our postural instructions, (2) head rotations would have been too slow to generate the 70 ms responses observed in the ear muscles, and (3) analysis of sternocleidomastoid EMG suggested that movements of the neck were rare, small, and unsystematic. Having documented the existence of directionally-appropriate responses of the ear muscles to brief novel sounds, we turn now to a qualitatively distinct type of attention.

### Experiment 2 – Endogenous Attention

To examine voluntary, goal-directed attention we used the classic, dichotic-listening paradigm (Cherry, 1953; Hillyard, Hink, Schwent, & Picton, 1973). Competing short stories were played over the two front or two back speakers. To increase motivation participants were allowed to choose, after a brief introduction, which of the two stories (podcasts) they would like to listen to. They were then told which speaker that story would be presented from. Our subjects were instructed to listen carefully while looking at a fixation cross and, as in the immediately preceding study, holding their head still on a chin rest. Upon completion, a new story was picked and the listening direction was switched to one of the other speakers. Recording methods were identical to those of the exogenous experiment. Muscle activity was quantified as mean, absolute EMG energy over the entire course of each 5 min listening trial.

As in the exogenous study, EMG energy at PAM and AAM was largest on the side to which attention was focused [Fig. 3; F(1, 19) = 15.2 and 4.6, respectively; *p* = 0.001 and 0.04; 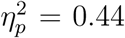 and 0.20]; an effect that is particularly strong in narrow–band middle frequency components of the signal, see Experiment 2 - figure supplement and 3.

A different pattern emerged for the other two muscles. Whereas TAM but not SAM activity had reflected lateralization of transient, exogenous attention, the reverse was true for sustained, goal-directed attention. That is to say, mean EMG energy at SAM was larger at the ipsi-than contralateral ear [F(1, 19) = 16.3; *p* = 0.001; 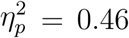] in Experiment 2, but there was no such difference for TAM. Another main effect indicated that activation of all four muscles was enhanced or at least tended to be so when participants listened to one of the two speakers that were slightly behind as opposed to in front of them [PAM, AAM, TAM, SAM: F(1, 19) = 5.7, 3.1, 8.1, and 12.0, respectively; *p* = 0.03, 0.09, 0.01, and 0.003; 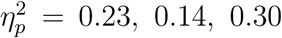, and 0.39]; data analogous to those of Fig 3, but for the front speakers, are provided in the Supplementary Information. These effects did not interact with each other or with age. Although PAM activity declines over time (Fig. 3), EMG energy of all three muscles is clearly sustained across the 5–min sessions. A corresponding sustained deflection of the pinna is also noticeable in a co–registered video, see `Experiment 2 - video supplement 1`.

As in experiment 1, we used the EOG and sternocleidomastoid EMG to rule out a systematic influence of gaze and head orienting, see Experiment 2 - figure supplement 3-6 and Experiment 2 - table supplement 1-5.

If sustained contraction of the auricular muscles is associated with a sense of fatigue, the expression “to strain one’s ears”, found in multiple languages, might be more than a metaphor (see also `Experiment 2 - video supplement 1`). The sustained character of pinna orienting suggests that optical or electrical measures might be combined non-redundantly with other objective, real-time measures of attentional focus. “Ear trackers” would complement eye trackers by offering a panoramic rather than narrow frontal range. They would complement electroencephalographic measures (de Cheveigné et al., 2018; Schäfer, Corona-Strauss, Hannemann, Hillyard, & Strauss, 2018) in that their sensitivity is exclusively spatial, rather than reflecting a context-specific mixture of modality, feature, location, and object representations. Moreover, EMG promises a decoding in near real–time (range of hundreds of milliseconds) and would not require information regarding the exogenous sound source.

## Discussion

These data provide compelling evidence that our brains retain vestigial circuitry for orienting the pinnae during both exogenous and endogenous modes of attention. The neural drive to our ear muscles is so weak that the movements are miniscule (see co– registered video data in the Supplementary Information) compared to those generated during broad smiles or voluntary ear wiggling. To understand what remains of the vestigial pinna–orienting system so as to exploit it for practical and scientific purposes it is helpful to take a comparative, phylogenetic approach (Hackley, 2015; Hackley, Ren, Underwood, & Valle-Inclán, 2017).

The ability to swivel and point the pinnae seems to have been lost during the transition from the primarily nocturnal lifestyles of prosimians to the diurnal ones of new world monkeys (Coleman & Ross, 2004). Mobility continued to decline as the ears became shorter and more rigid (Coleman & Ross, 2004; Waller, Parr, Gothard, Burrows, & Fuglevand, 2008). The musculature degenerated. For example, an inferior auricular muscle to oppose SAM still exists in lesser apes (gibbons and siamangs) (Burrows, Diogo, Waller, Bonar, & Liebak, 2011), but not in chimpanzees (Burrows et al., 2011) or humans (Cattaneo & Pavesi, 2014). The use of proprioceptive information to adjust auditory processing in accordance with pinna position, orientation, and shape is known to occur in cats (Kanold & Young, 2009). This might be possible in old world monkeys, which have muscle spindles (Lovell, Sutton, & Lindeman, 1977), but not in humans, whose auricular muscles lack these structures (Cattaneo & Pavesi, 2014).

Our auriculomotor control system (which might include the superior colliculi (Stein & Clamann, 1981; Henkel & Edwards, 1978), the paralemniscal zone (Henkel & Edwards, 1978), and a human homolog of the macaque premotor ear-eye field (Lanzilotto, Perciavalle, & Lucchetti, 2013)) apparently receives little or no proprioceptive feedback. Cutaneous rather than proprioceptive afference may be involved when one feels an ear twitch upon registering the faint sound of a door opening from behind^1^. The present data suggest that such phenomena are not necessarily a product of the imagination. They may reflect a response by our ancient pinna-orienting system.

## Methods

### Participants

Both older (N = 12, M = 62.7 ±5.9 y, 8 F, all right-handed) and younger adults (N = 16, M = 24.1 ± 3.1 y, 8 F, 15 right-handed, 1 left-handed) volunteers in Experiment 1 had age-typical, pure tone audiometric thresholds (1, 2, 4, and 8 kHz; young < 20 dB; old < 40 dB). After the 8th participant, Experiment 2 was altered in several ways (e.g., four stimulus directions rather than two). Only data from the final 21 subjects were retained for Experiment 2. The two groups in this experiment comprised 11 older adults (M = 62.6 ± 6.2 y, 8 F, all right-handed) and 10 younger adult (M = 24.1 ± 3.6 y, 5 F, 9 right-handed, 1 left-handed). The study was approved by the responsible ethics committee (ethics commission at the Ä rztekammer des Saarlandes, Saarbrücken, Germany).

### Stimuli and Tasks

The four active loudspeakers (KH120A, Neumann, Germany) were at head level, 115 cm. Sounds in Experiment 1 and 2 were reproduced with a soundcard (Scarlett 18i20, Focusrite, UK). The experimental paradigms were programmed using software for scientific computing (Matlab, Mathworks, USA) and Psychtoolbox 3. In Experiment 1, sounds lasted 1.7 – 10.0 s, were delivered every 15 – 40 s, and had an average intensity of 70 dBLc, except for foot steps (65 dBLc). Each of the nine stimuli (lemur howling, dog barking, helicopter flying, cell phone vibrating, birds singing, baby crying, mosquito buzzing, foot steps, and traffic jam) was repeated four times (i.e., once per speaker).

In Experiment 2, the stories were 5 min long, with an average intensity of 50 dBLa for younger and 60 dBLa for older participants. Participants answered content questions at the conclusion of each condition in this experiment.

### Electrophysiological Recordings & Signal Processing

Surface EMGs were recorded with non-recessed, Ag/AgCl electrodes (BME4, BioMed Electrodes, USA), which were 4 mm in diameter for TAM and 6 mm (BME6) in all other cases. The signals were AD-converted at 9600 Hz and 24 bit resolution per channel (4 × USBamp, g.tec GmbH, Austria). Skin temperature, skin resistance, electrocardiograms, and EOGs were also recorded. Preliminary tests involving Wilson’s phenomenon and ear wiggling indicated that EMG from TAM electrodes was not an artifact of volume conduction from PAM or SAM. Sternocleidomastoid EMG signals were zero–phase bandpass filtered from 60 to 100 Hz (FIR, 2000th order), the auricular EMG signals from 10 to 1000 Hz (FIR, 2000th order) with a notch filter at 50 Hz (IIR, 2nd order). HEOG signals were zero-phase filtered from 0.01 to 20 Hz (IIR, 2nd order). All signals were then downsampled to 2400 Hz for further processing. The statistical analysis was performed using repeated measures ANOVA (with IBM SPSS Statistics 26). Within–subjects factors were stimulus–muscle correspondence (ipsi-vs. contralateral responses) and anteriority (front or back speakers). The only between– subjects factor was age (older vs. younger subjects). All main and interaction effects were tested.

#### Exogeneous (transient) data

Root-mean-square (RMS) envelopes of the filtered and downsampled EMG signals were calculated with a sliding window of 150 samples (62.5 ms). The data were then segmented into epochs extending from 3 s prior to stimulus onset until 3 s following termination of the auditory stimulus, which was of variable duration. Epochs were baseline corrected with respect to the mean RMS envelope amplitude of the pre–stimulus interval. Amplitudes were normalized separately for each participant and muscle according to the largest value among the four stimulus presentations at any time point within the epoch data. Each participant’s data were pooled according to whether the side of the stimulus and recorded muscle did or did not match, and then were averaged into contralateral and ipsilateral waveforms. Eighteen trials contributed to each of these per subject waveforms, 9 from the left speaker and 9 from the right. Mean amplitudes were computed across a measurement window extending from 100 to 1500 ms following stimulus onset and were then subjected to statistical analysis.

#### Endogenous (sustained) data

Artifacts of the filtered and downsampled EMG data were reduced by averaging the signal energy of 1 s, non-overlapping segments and rejecting segments that deviated by more than two standard deviations. For each participant, the mean energy of a given channel during the four listening conditions (left/right × front/back) was calculated and then normalized to the largest value across the 5–min run. These normalized data were then averaged into ipsilateral/contralateral categories and subjected to statistical analysis.

#### EMG Time–Frequency Decomposition (for Experiment 2 – figure supplement 2 and 3)

As the rather sustained muscle activity during endogenous attention might be reflected in low frequency components according to convolution models of EMG, filtered and downsampled EMG signals in Experiment 2 were decomposed into eight frequency bands by a nonsubsampled octave–band filter bank (5th order Daubechies filter). Each frequency band was then further processed in the same fashion as the broadband signals reported in the main text.

### Computer Vision Setup and Motion Analysis (for the video supplements in Experiment 1 and 2)

Videos were acquired using four Ximea MQ022CG–CM color sensors with a resolution 1936 × 1216 at 120 frames per second and an exposure of 2 ms. Two cameras were positioned on each side of the head and focused on the ears to record pairwise stereo videos. We used hardware triggering for all four cameras and recorded each camera onto a separate m.2 solid–state–drive to reduce frame loss. We used a KOWA 35 mm macro lens with an aperture of F0.4 which gave us a close–up view of the ear with acceptable depth of field to allow slight movements towards the camera and enough distance such that the cameras did not cast shadows on to the scene. We illuminated the face uniformly with flicker–free LED studio illumination. The cameras were calibrated with the stereo camera calibrator app from Mathworks MATLAB Computer Vision System Toolbox. Calibration was performed whenever camera adjustment required re– alignment of relative stereo camera positions or a change of focus of one of the cameras. For 3D reconstruction and motion visualization/quantification, we used functions from the Mathworks MATLAB Computer Vision System Toolbox and custom written code. Our analysis system was able to reduce redundancies in optic flow and stereo depth estimation by exploiting the unilateral scene composition and limited degrees of freedom for ear and head movements. For 3D reconstructions, we initialized a sequence with one initial estimation of disparity and subsequently tracked points independently for the left and right image sequence.

We tracked points with respect to the first frame of the sequence as reference frame with dense optical flow initialized with a rigid motion estimation. Motion was visualized with a Lagrangian motion magnification approach that had a constant magnification factor with respect to the reference frame and prior removal of affine motion with respect to manually selected stable points. The results of the motion analysis with and without magnification can be seen in the supplementary videos to Experiment 1.

## Supporting information

Exp1VideoSupp2

Exp2VideoSupp1

Exp1VideoSupp1

## Acknowledgements

This study was partially supported by the German Federal Ministry of Education and Research, Grant No. BMBF-FZ 03FH004IX5 (PI: D.J.S). We thank Larissa Arand for assistance with data collection and Becca Sullinger for the artwork in Figure 1.

## Author Information

Daniel J. Strauss, Farah I. Corona-Strauss, Andreas Schroeer, Philipp Flotho are with the Systems Neuroscience & Neurotechnology Unit, Faculty of Medicine, Saarland University & htw saar, Homburg/Saar, Germany. Ronny Hannemann is with the Audiological Research Unit, Sivantos GmbH, Erlangen, Germany, and Steven A. Hackley is with the Department of Psychological Sciences, University of Missouri–Columbia, Columbia, MO, USA.

## Contributions

D.J.S., F.C.-S., R.H. and S.A.H. conceived and designed the study. F.C.-S. recruited participants and conducted the experiments with support of P.F. The data were processed and analysed by A.S., D.J.S., and P.F. The manuscript was written by D.J.S. and S.A.H. All authors approved the submitted version.

## Video and Figure Supplements for Experiment 1 and 2

### 1. Supplementary Videos & Video Legends

Three supplementary videos were uploaded.

#### Video 1 (filename: “Exp1VideoSupp1.mp4”)

Legend to video 1: `Experiment 1 - video supplement 1`: Ear movement example from a trial with a novel sound at the right posterior speaker. The right half of the display portrays evoked movements of the ipsilateral pinna in three ways. The large video clip of the pinna uses digital magnification to render the overall pattern of movement apparent. The color overlay in these videos indicates the motion magnitude. Just below the video and to the right, an unrectified EMG recording of the postauricular muscle is shown in co–registration with the video. The global head motion was reduced by a 2-dimensional rigid pre-registration with respect to a set of manually specified reference points on the head (see also Experiment 1—figure supplement 1, below). The 3-dimensional graph medial to the 2-D graph includes a vector that indicates moment–by–moment changes in EMG activity of the superior auricular muscle (SAM, the vertical axis), transverse auricular muscle (TAM, a horizontal axis), and the difference between activity in the posterior and anterior auricular muscles (PAM-AAM, the other horizontal axis). The left half of this video gives corresponding information for the contralateral ear which, consistent with evidence presented in the main text, was not as active as the ipsilateral one.

#### Video 2 (filename: “Exp1VideoSupp2.mp4”)

Legend to video 2: `Experiment - video supplement 2`: The right ear example from the video supplement before, but with 4 different videos in sequence. The first video of the sequence shows the raw recording (without digital magnification). The second video shows the digitally magnified motion, the third video shows the magnified motion with color overlay as in the video supplement before, and the fourth video shows the 3 dimensional motion from a different angle. This video sequence shows the impact of the digital motion magnification and the depth information about ear motion that can be derived from a stereo computer vision setup such as the one used here.

#### Video 3 (filename: “Exp2VideoSupp1.mp4”)

Legend to video 3: `Experiment - video supplement 1`: Ear movement example from a participant who exhibited exceptionally large, long–lasting involuntary auricular muscle activations and ear motion during the endogenous attention task in Experiment 2. The attention of the participant was directed to the story played from the posterior right speaker. The organization of the plots and co–registration is as in Experiment 1 - video supplement 1. However, this time the raw videos without digital magnification are shown. The raw videos are played faster, time–locked to the time axis given in minutes in the 1 dimensional plots of the rectified postauricular muscle activity. Note that time–axis reflects the entire timeline including the instructions and the introduction to the stories before the directional listening task. The listening task started at approximately 2min. The video also documents the end of the listening task (around 7min) accompanied with a time–locked offset of the muscle activation and pinna displacement. A causal relation of the rectified postauricular muscle activity and the motion magnitude in the videos is clearly noticeable, especially for the ipsilateral ear.

### 2. Supplementary Figures

Nine supplementary figures are shown below (one figure per page and accompanying captions).

**Experiment 1 - figure supplement 1:**
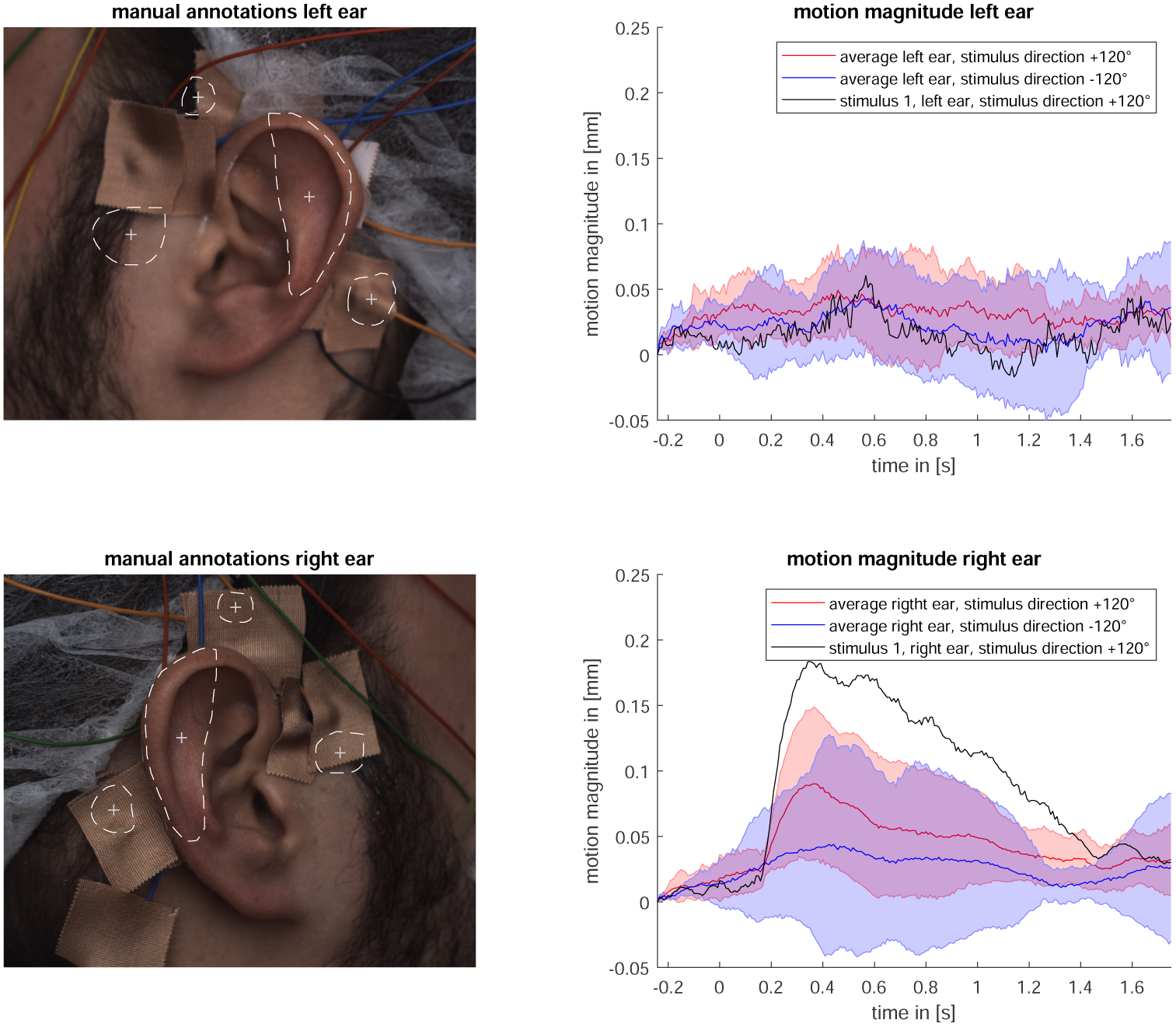
Analysis of video recordings from one participant who exhibited submillimeter movements in response to stimulation in Experiment 1 (Exogenous Attention). Given the time *t*^*i*^ of a stimulus, we analysed 2 s video segments with pre–stimulus onset 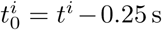. The frames at 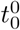 of the first stimulus (direction +120°) of the left and right recordings were manually annotated with four regions of interest (ROI) (left images). The centroid *P*_0_ of the ROI located on the ear was used to track the pinna motion and the three centroids 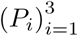 of the ROIs around the ears were used to define a basis *B*′ of a coordinate system where the coordinates of points on the ear are invariant under head motion. Each point inside the ROIs sampled as in the first frame was projected to the respective right camera frames and forward-warped with respect to the motion to the reference frame for all cameras. The motion magnitude is given as the norm of *P*_0_ at time *t*^*i*^ with respect to the basis *B*′ referenced to *P*_0_ at 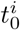. On the right are the time courses of the mean motion magnitude and standard deviation over the stimuli (±120°) for the left and the right ear as well as the time course of the first stimulus from the right back speaker. This is the same trial that is portrayed in video 1 and video 2. Due to a large head rotation during the recording, the first −120° stimulus was excluded from the evaluation.

**Experiment 1 - figure supplement 2:**
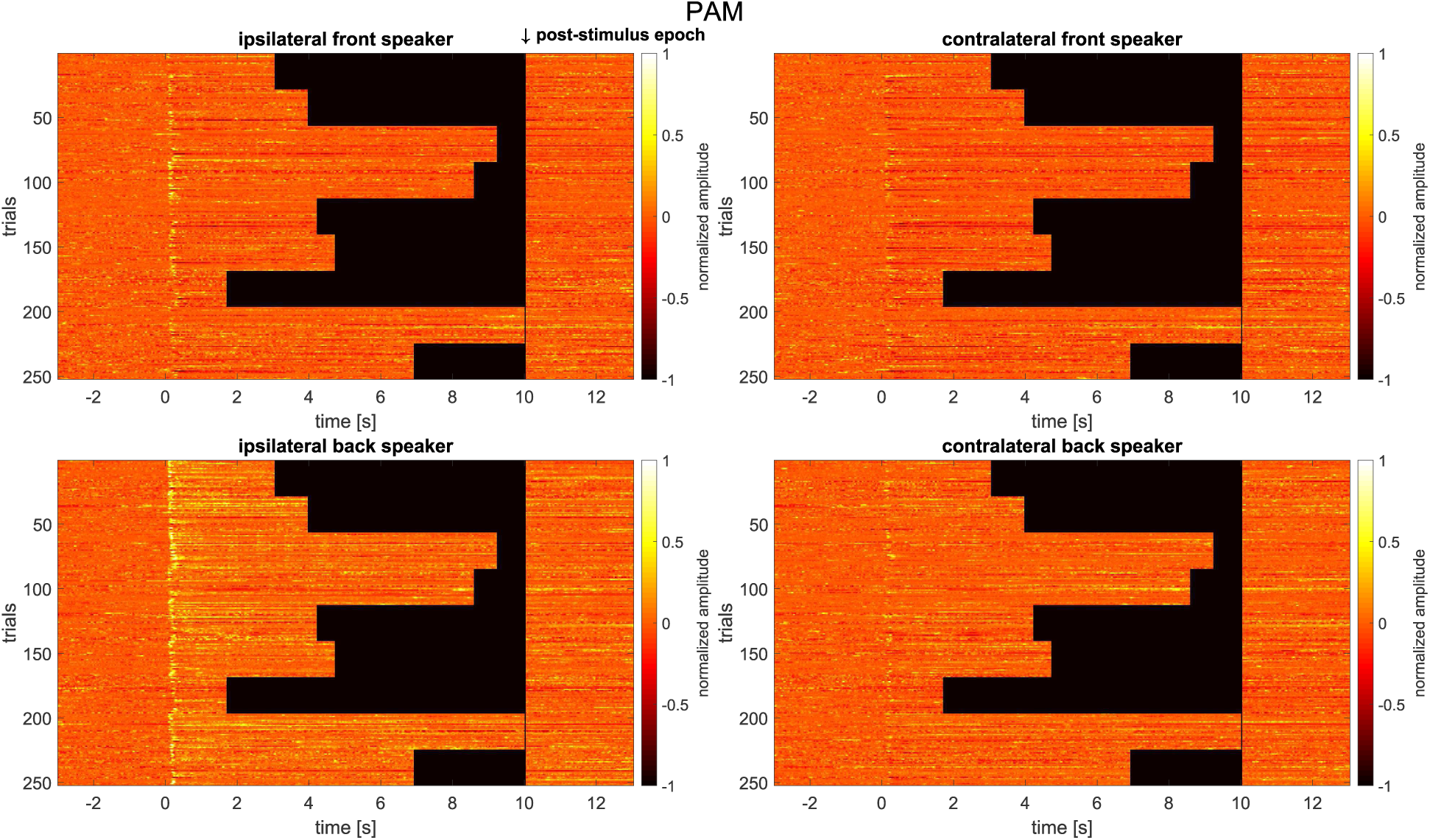
Individual epoch matrix of the baseline corrected and normalized event-related PAM electromyograms for the recording ipsilateral (left panel) and contralateral (right panel) to stimulation; top: front speakers (30°), bottom: back speakers (120°). Shown are 252 trials in total (*N* = 28 participants × 9 stimuli) per direction. The amplitude of the normalized data is encoded in colors (white/yellow: large; dark orange: small). Individual epochs are grouped according to stimulus duration, with 28 participants stacked to the left of the black area. It is notable that the individual epochs (i.e, one row in the matrix) show a rather consistent response (with an onset of about 70 ms) across the 9 stimuli/28 participants per direction/speaker in contrast to the 2 s period immediately following stimulus offset (shown separately on the right of the individual plots).

**Experiment 1 - figure supplement 3:**
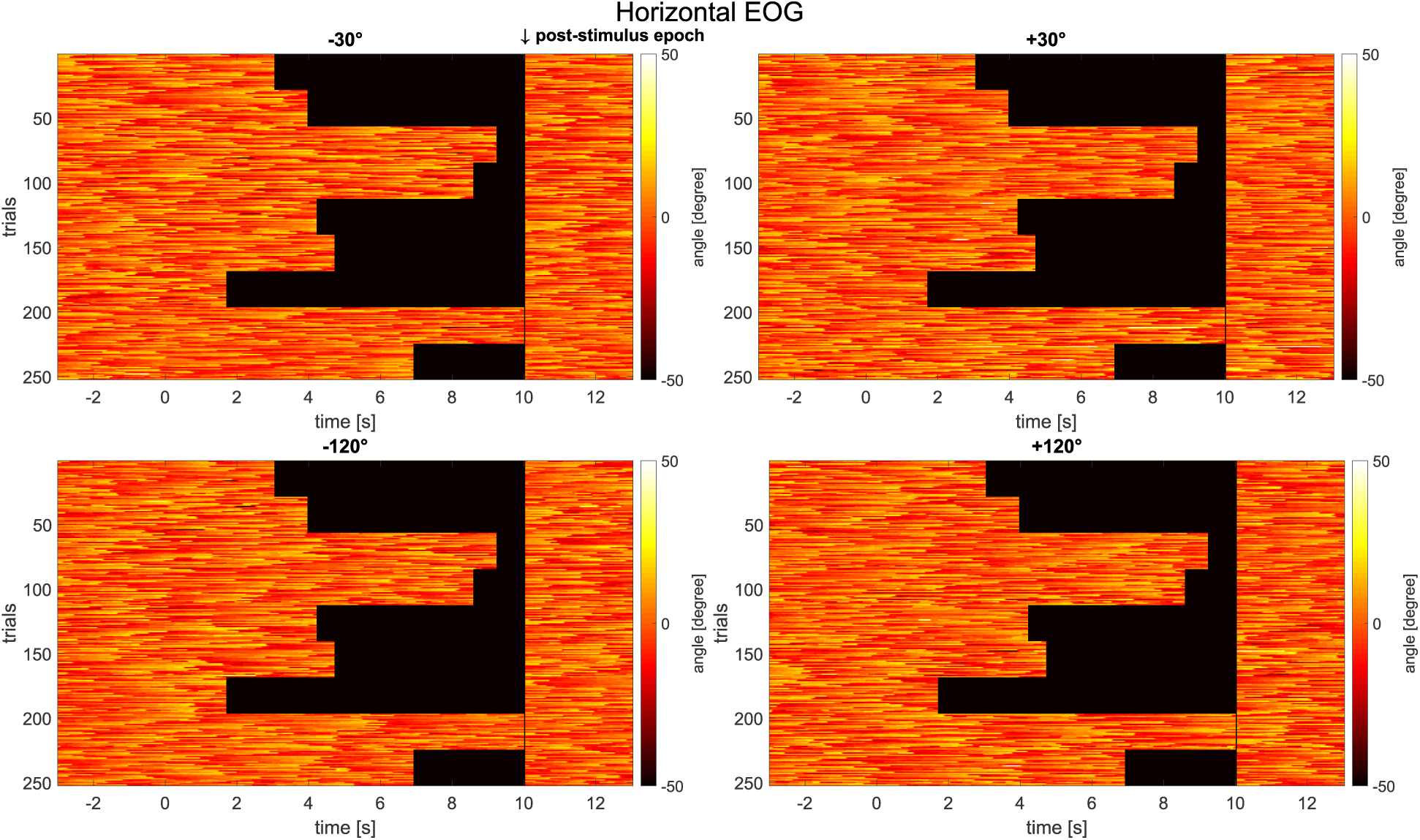
Individual epoch matrix of the event–related horizontal EOG. Each matrix represents the responses to stimuli from a certain direction. Shown are 252 trials in total (*N* = 28 participants × 9 stimuli) per direction. The angle of the horizontal EOG is encoded in colors. A positive angle indicates a right gaze direction. Individual epochs are grouped according to stimulus duration, with 28 participants stacked to the left of the black area. In contrast to the EMG data shown in the previous figure, these eye–movement recordings show no consistent, stimulus-locked response. This supports our claim that pinna responses to novel stimuli are not secondary to the oculo–auricular phenomenon.

**Experiment 1 - figure supplement 4:**
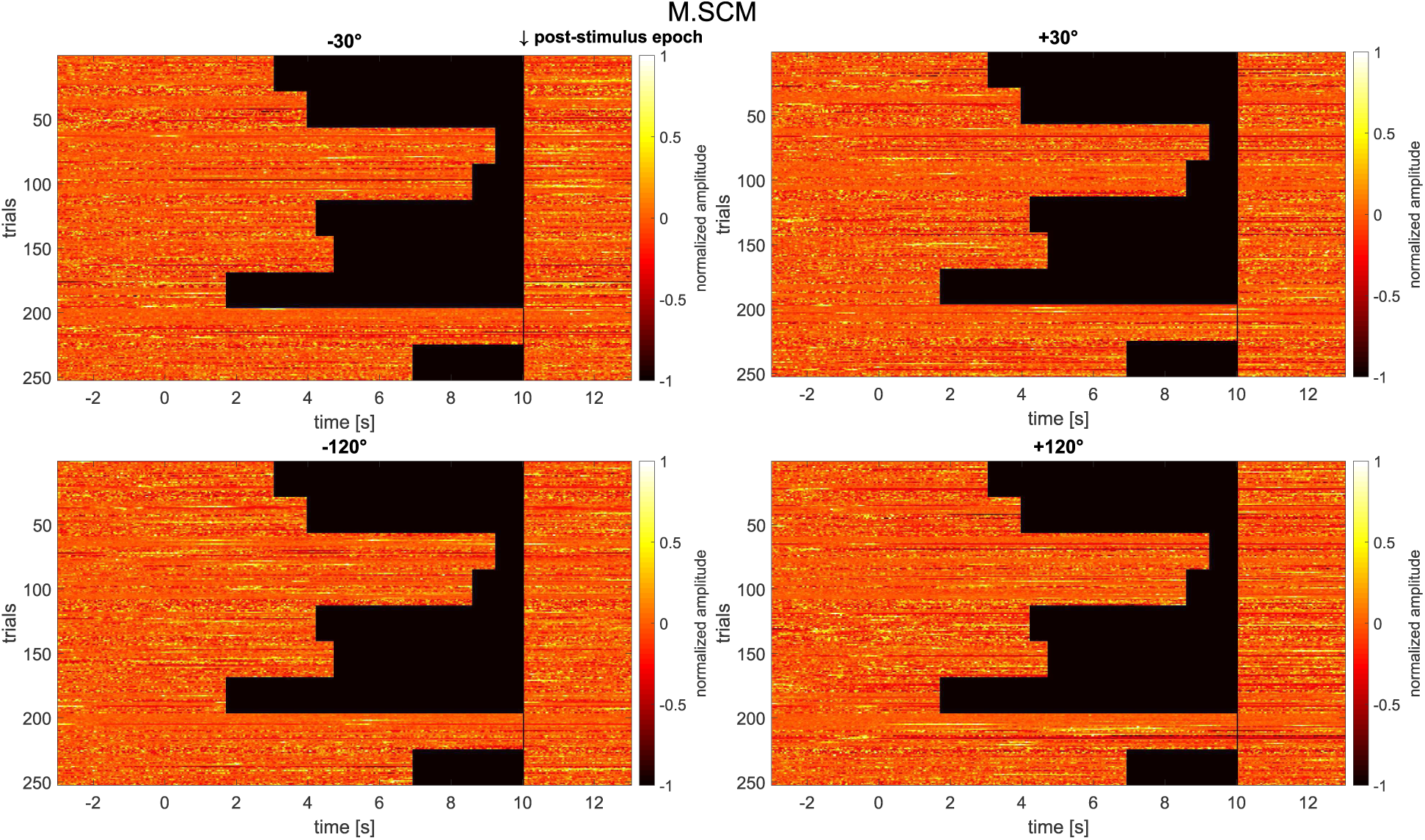
Individual epoch matrix of the baseline corrected and normalized event-related electromyograms recorded at the M. sternocleidomastoideus. Each matrix represents the responses to stimuli from a certain direction. Shown are 252 trials in total (*N* = 28 participants × 9 stimuli) per direction. As in the previous two figures, increases in amplitude are encoded as lighter colors. Individual epochs are grouped according to stimulus duration, with 28 participants stacked to the left of the black area. In contrast to the EMG data shown earlier for the PAM muscle, these neck–muscle recordings show no consistent, stimulus–locked responses. This supports our claim that pinna responses to novel stimuli are not secondary to the vestibulo–auricular reflex, if the latter exists in our species.

**Experiment 1 - figure supplement 5:**
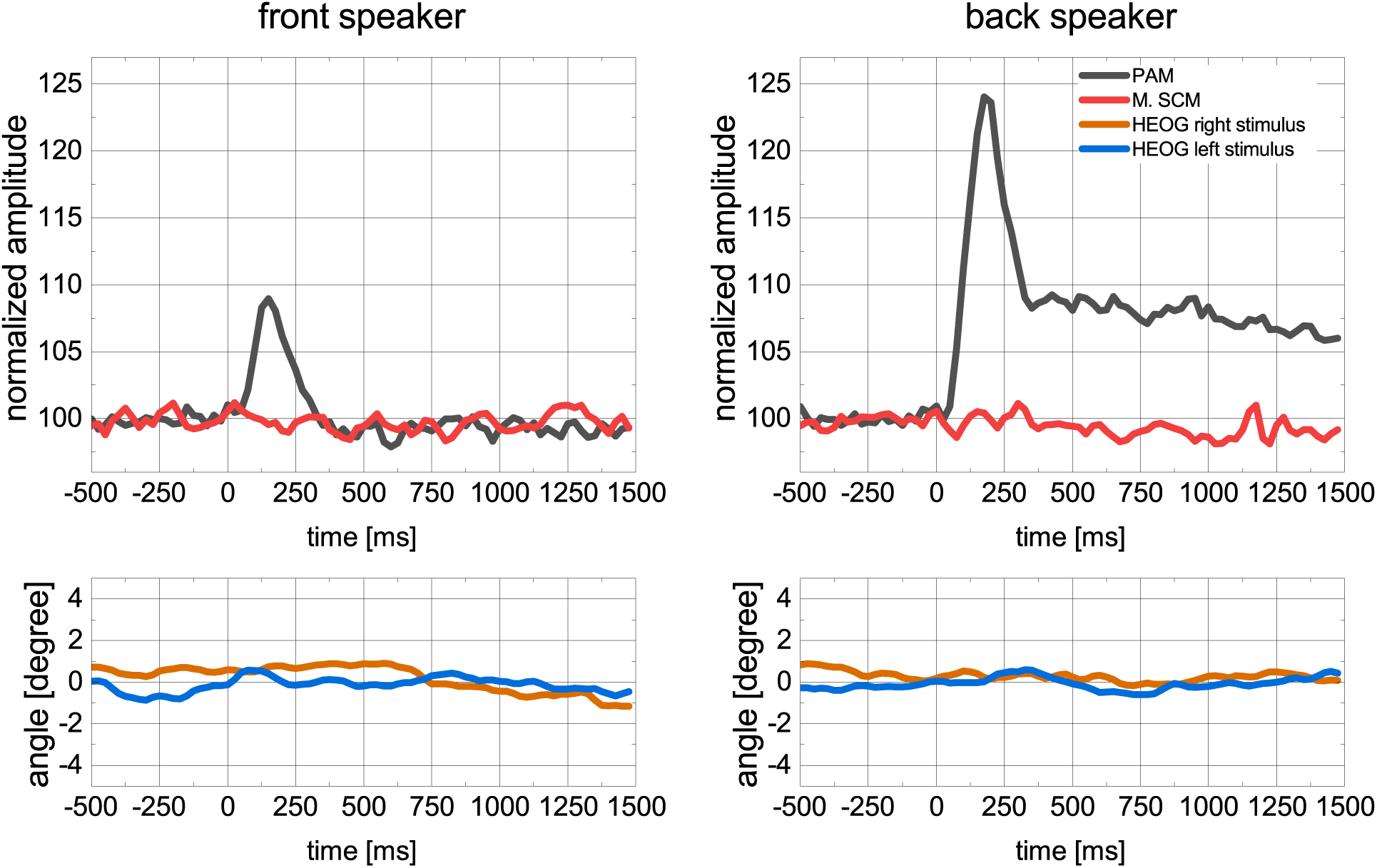
Grand average of the baseline corrected and normalized electromyograms recorded at the ipsilateral PAM, M. sternocleidomastoideus (M.SCM) and horizontal EOG (HEOG) across all subjects and stimuli. A positive deflection in the HEOG indicates a gaze shift in the direction of the stimulus. Note that in contrast to the PAM, there is no stimulus locked activity in either M.SCM or HEOG.

**Experiment 2 - figure supplement 1:**
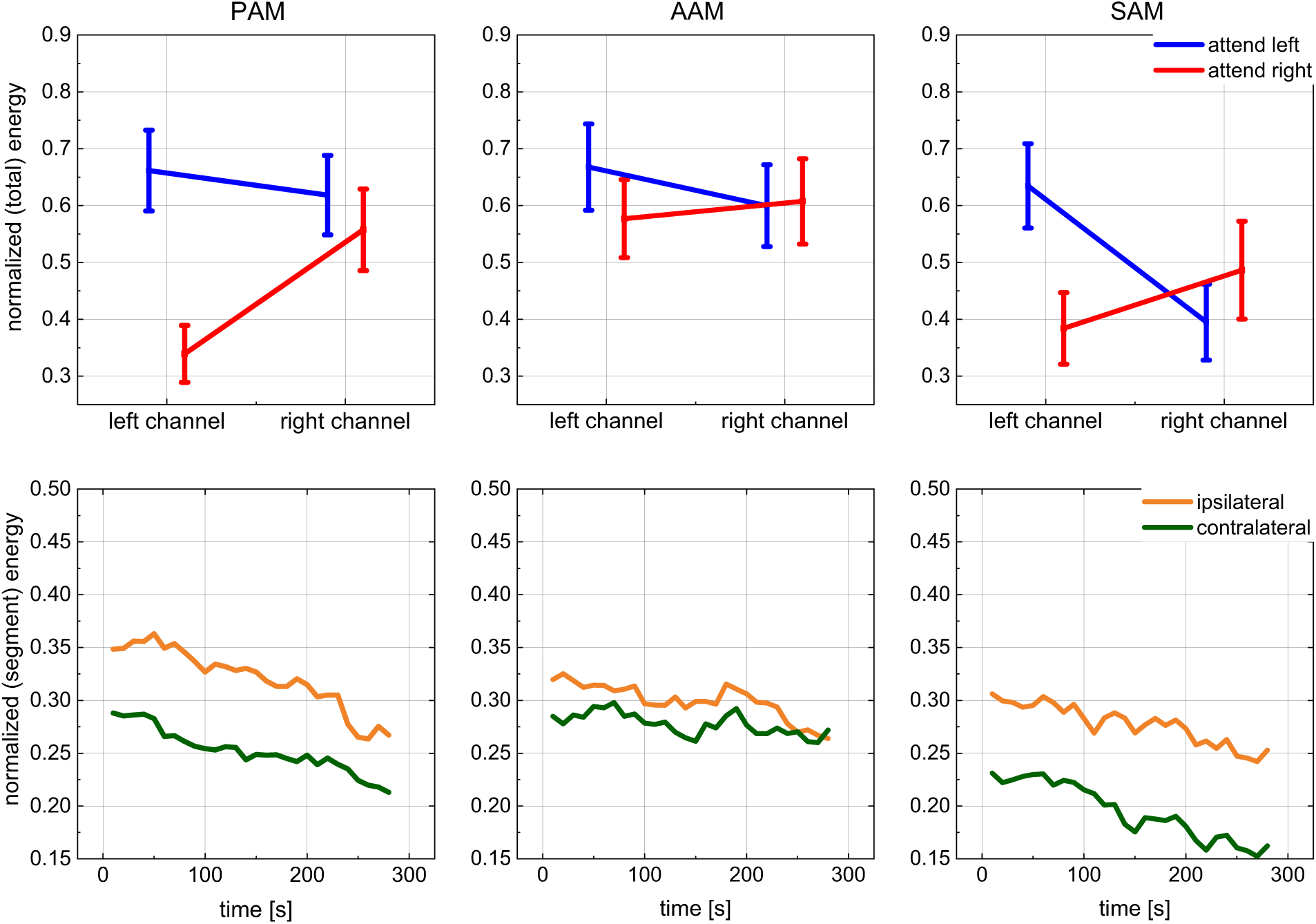
These graphs are analogous to those of Fig. 3 in the main text but are for the front speakers: Grand average of PAM, AAM, and SAM activity when stories were played from the front speakers (±30°). Top: normalized total energy of the left/right recording channels (broadband electromyograms) during attention to the left or right story (bars represent the standard error). Bottom: time–resolved activity after pooling the ipsi– and contralateral signals with a segment–wise normalization. Each sampling point represents the energy induced in consecutive 10 s segments. It is notable that the described effect is less strong for the stimulation from the front as compared to back stimulation. This is to be expected for an attention effect, given that the front speakers were much closer together than the rear speakers.

**Experiment 2 - figure supplement 2:**
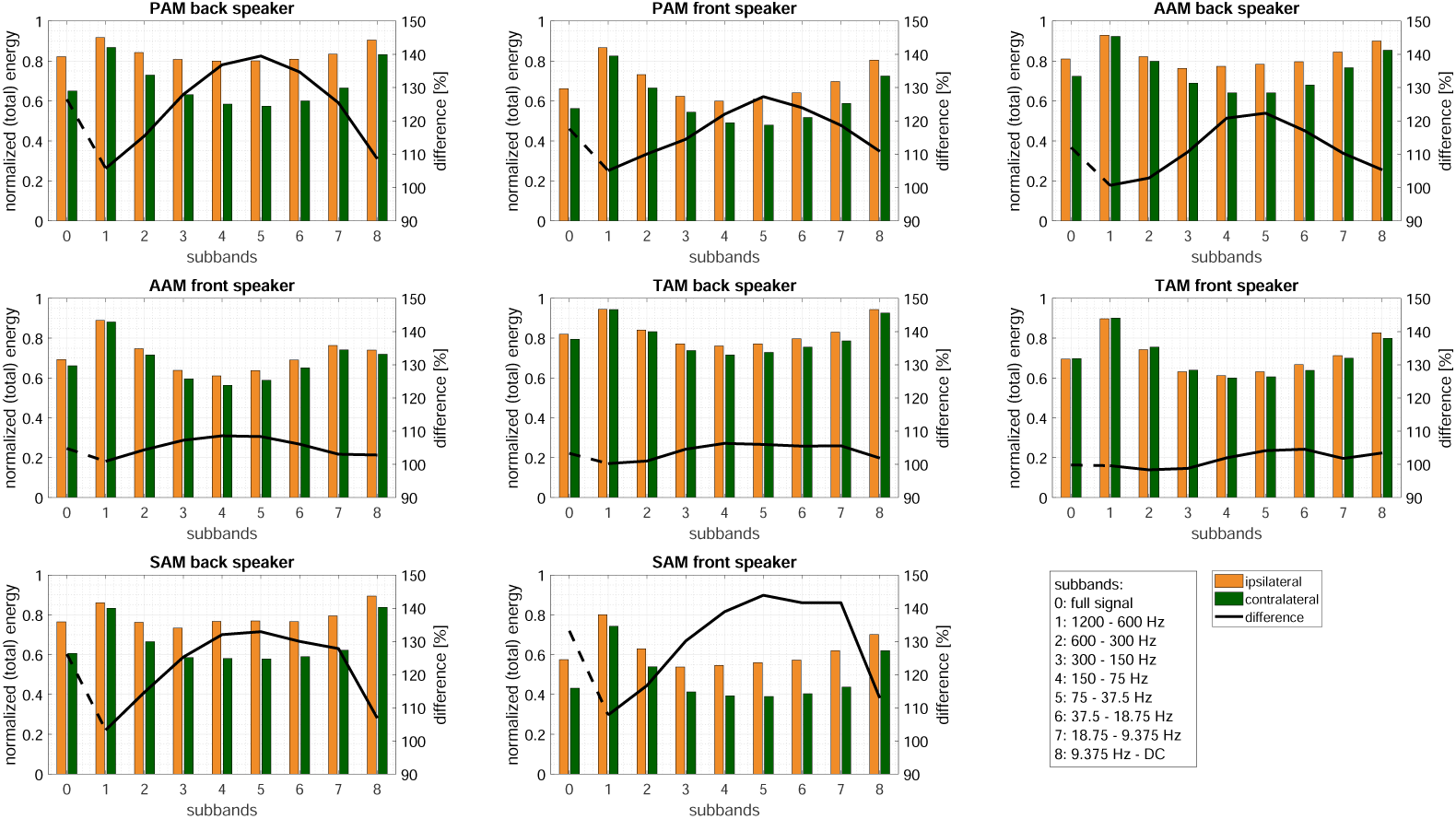
Grand average of the PAM, AAM, and SAM activity when stories were played from front (±30°) and back speakers (±120°). The normalized total energy in each frequency subband is plotted for the left/right recording channels during attention to the left or right story, along with their difference (black line). It is notable that the lowest and highest frequency bands contribute little to the difference. In fact, the middle bands 5 and 6 show the largest attention effect (see also Experiment 2 - figure supplement 3). Technical applications such as attentionally controlled hearing might make use of this finding.

**Experiment 2 - figure supplement 3:**
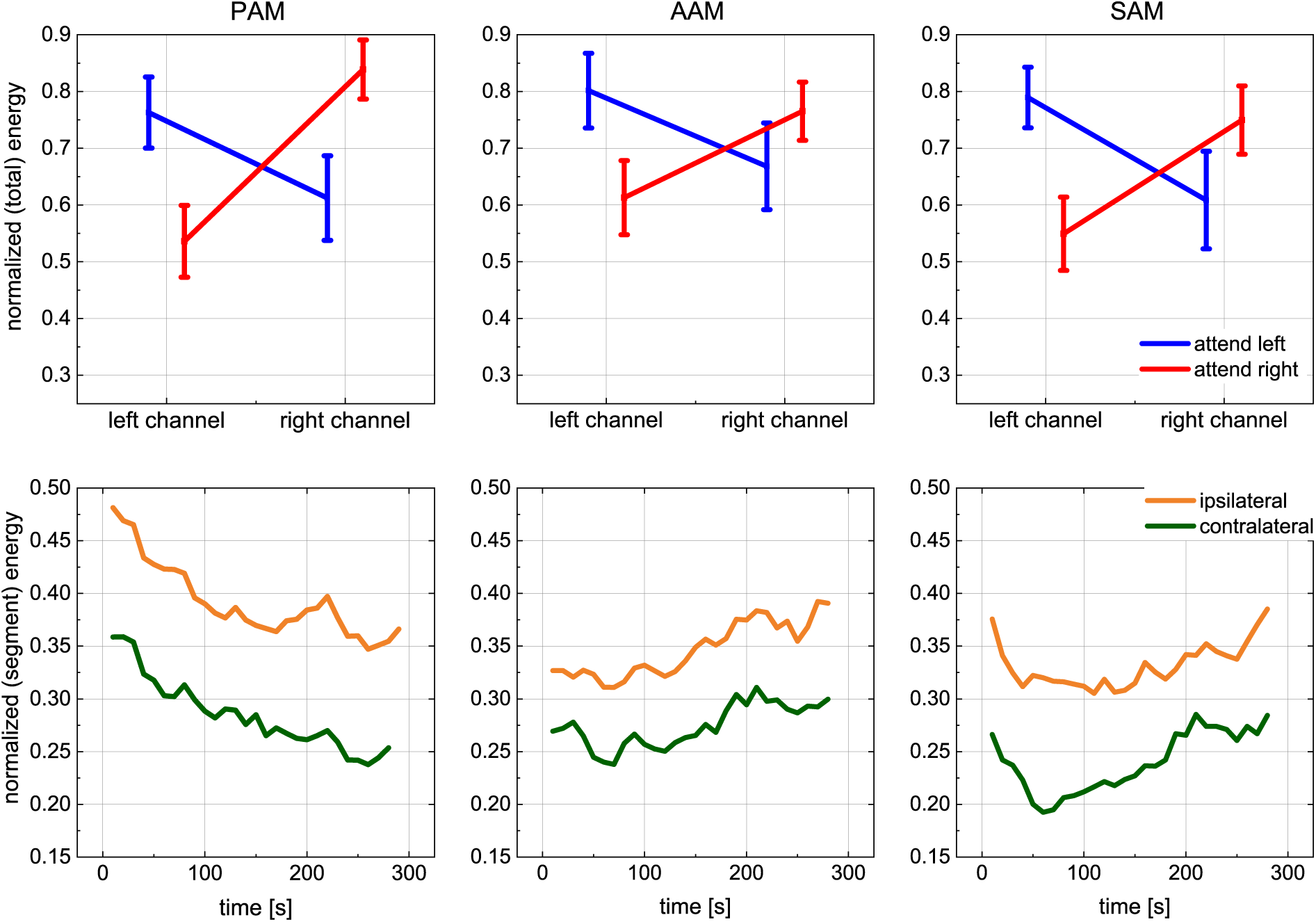
Analogous to Fig. 3 in the main text but for frequency band 5 (75 – 150 Hz): Grand average of the PAM, AAM, and SAM activity when stories were played from the front speakers (±120°). Top: normalized total energy of the left/right recording channels when attending to the left or right story (bars represent the standard error). A significant interaction of recording channel and attention direction was observed [PAM, AAM, TAM, SAM: F(1, 19) = 17.2, 8.2, 0.5, and 24.0, respectively; *p* = 0.001, 0.01, 0.51, and 0.001; 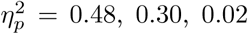, and 0.56]; Bottom: time–resolved activity after pooling the ipsi– and contralateral signals with a segment–wise normalization. Each sampling point represents the energy induced in consecutive 10 s segments.

**Experiment 2 - figure supplement 4:**
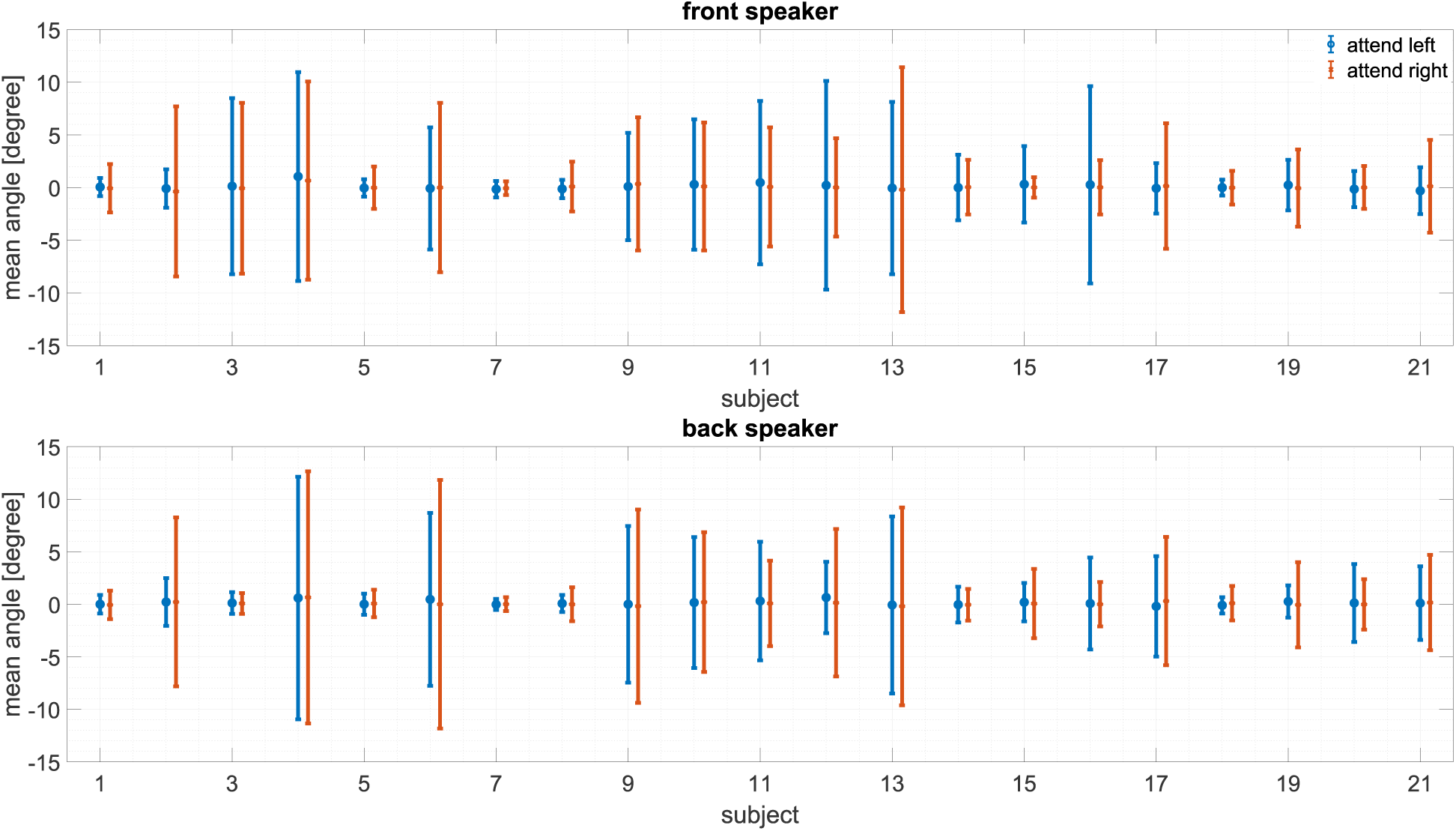
Mean angle and standard deviation of the horizontal EOG deflections for each individual subject; top: attending front speakers (±30°), bottom: attending back speakers (±120°). A positive deflection in the HEOG indicates a gaze shift in the direction of the stimulus. While some subjects display a large standard deviation, the mean angle during the measurements is always well below 5°, indicating no bias to the attended direction. These data further support our claim that directionally–specific activity in the auricular muscles is not secondary to Wilson’s (1908) oculo–auricular phenomenon.

### 3. Supplementary Tables

The following tables summarize the statistical results when using gaze shifts and the neck muscle activity additionally for the artifact rejection in the individual muscles electromyograms. The results demonstrate that the inclusion of this information has no major impact, strengthening the conclusions made from Experiment 2 – figure supplement 3-6.

**Table 1: Experiment 2 - table supplement 1:**
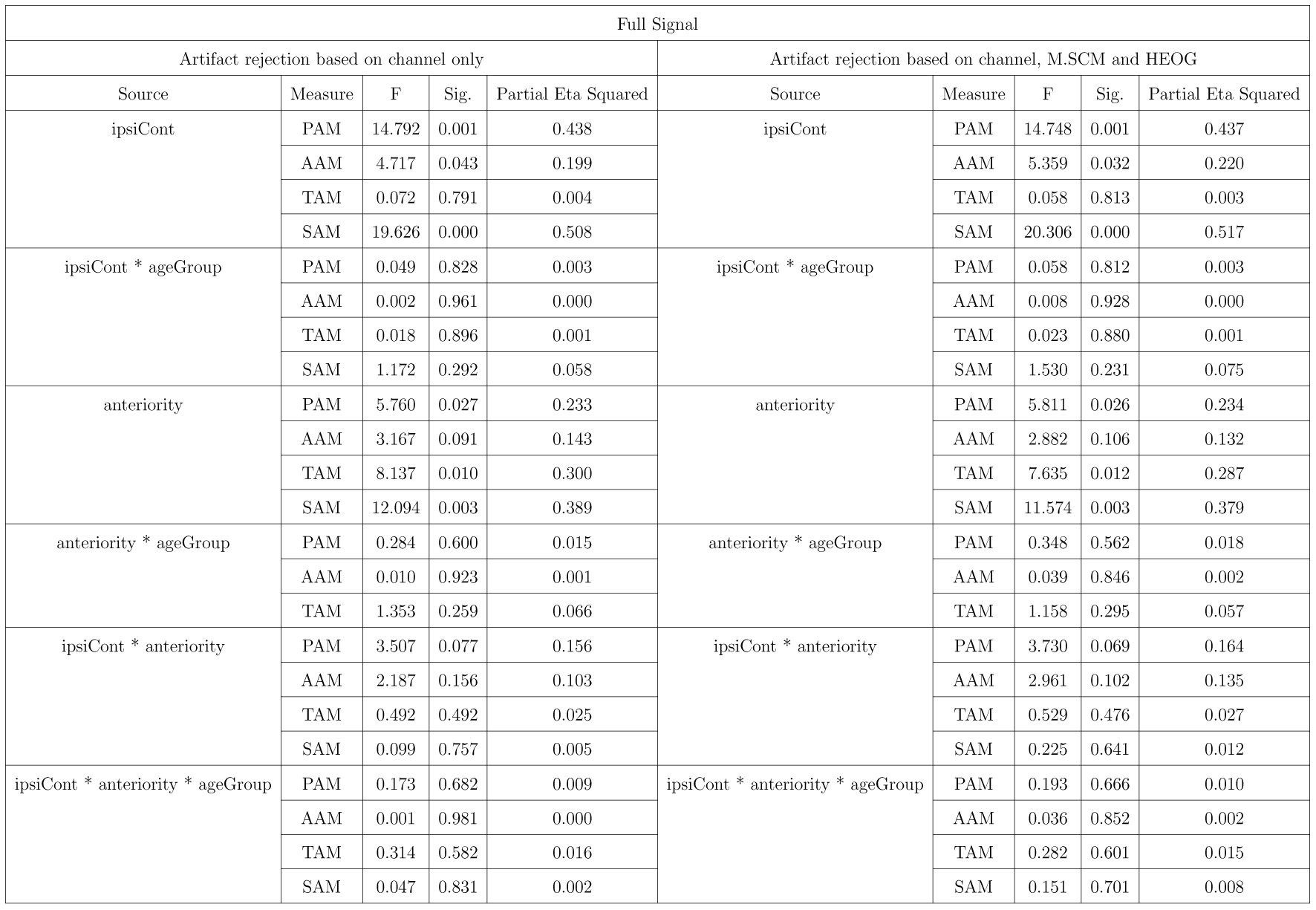
Results of the repeated measures ANOVA with three witin-subjects factors for all four auricular muscles. ipsiCont: ipsilateral vs. contralateral channel (with respect to the attended side). ageGroup: older vs. younger adults. anteriority: stimulus presentation from the front or back speakers. The left columns show the results when only the corresponding channel was used to reject artifacts, while the right columns display the results when artifact rejection additionally considers the M.SCM and HEOG.

**Table 2: Experiment 2 - table supplement 2:** Results of the repeated measures ANOVA with three witin-subjects factors for the frequency decomposed PAM. f1 to f8 refer to the subbands in descending order of frequency. ipsiCont: ipsilateral vs. contralateral channel (with respect to the attended side). ageGroup: older vs. younger adults. anteriority: stimulus presentation from the front or back speakers. The left columns show the results when only the corresponding channel was used to reject artifacts, while the right columns display the results when artifact rejection additionally considers the M.SCM and HEOG.

| Subbands PAM |  |  |  |  |  |  |  |  |  |
| --- | --- | --- | --- | --- | --- | --- | --- | --- | --- |
| Artifact rejection based on channel only |  |  |  |  | Artifact rejection based on channel, M.SCM and HEOG |  |  |  |  |
| Source | Measure | F | Sig. | Partial Eta Squared | Source | Measure | F | Sig. | Partial Eta Squared |
| ipsiCont | f1 | 8.492 | 0.009 | 0.309 | ipsiCont | f1 | 8.282 | 0.010 | 0.304 |
|  | f2 | 10.668 | 0.004 | 0.360 |  | f2 | 10.735 | 0.004 | 0.361 |
|  | f3 | 12.544 | 0.002 | 0.398 |  | f3 | 12.282 | 0.002 | 0.393 |
|  | f4 | 14.650 | 0.001 | 0.435 |  | f4 | 14.413 | 0.001 | 0.431 |
|  | f5 | 17.185 | 0.001 | 0.475 |  | f5 | 17.086 | 0.001 | 0.473 |
|  | f6 | 16.865 | 0.001 | 0.470 |  | f6 | 17.133 | 0.001 | 0.474 |
|  | f7 | 17.671 | 0.000 | 0.482 |  | f7 | 18.116 | 0.000 | 0.488 |
|  | f8 | 10.320 | 0.005 | 0.352 |  | f8 | 10.432 | 0.004 | 0.354 |
| ipsiCont * ageGroup | f1 | 0.125 | 0.728 | 0.007 | ipsiCont * ageGroup | f1 | 0.158 | 0.695 | 0.008 |
|  | f2 | 0.368 | 0.551 | 0.019 |  | f2 | 0.418 | 0.525 | 0.022 |
|  | f3 | 0.077 | 0.785 | 0.004 |  | f3 | 0.107 | 0.747 | 0.006 |
|  | f4 | 0.003 | 0.959 | 0.000 |  | f4 | 0.001 | 0.982 | 0.000 |
|  | f5 | 0.000 | 0.985 | 0.000 |  | f5 | 0.000 | 0.992 | 0.000 |
|  | f6 | 0.013 | 0.911 | 0.001 |  | f6 | 0.013 | 0.909 | 0.001 |
|  | f7 | 0.004 | 0.950 | 0.000 |  | f7 | 0.007 | 0.936 | 0.000 |
|  | f8 | 0.000 | 0.985 | 0.000 |  | f8 | 0.003 | 0.958 | 0.000 |
| anteriority | f1 | 2.409 | 0.137 | 0.113 | anteriority | f1 | 2.430 | 0.136 | 0.113 |
|  | f2 | 4.083 | 0.058 | 0.177 |  | f2 | 4.059 | 0.058 | 0.176 |
|  | f3 | 6.217 | 0.022 | 0.247 |  | f3 | 6.269 | 0.022 | 0.248 |
|  | f4 | 6.403 | 0.020 | 0.252 |  | f4 | 6.521 | 0.019 | 0.256 |
|  | f5 | 6.260 | 0.022 | 0.248 |  | f5 | 6.362 | 0.021 | 0.251 |
|  | f6 | 5.275 | 0.033 | 0.217 |  | f6 | 5.289 | 0.033 | 0.218 |
|  | f7 | 4.308 | 0.052 | 0.185 |  | f7 | 4.349 | 0.051 | 0.186 |
|  | f8 | 11.086 | 0.004 | 0.368 |  | f8 | 11.072 | 0.004 | 0.368 |
| anteriority * ageGroup | f1 | 1.983 | 0.175 | 0.095 | anteriority * ageGroup | f1 | 2.097 | 0.164 | 0.099 |
|  | f2 | 1.137 | 0.300 | 0.056 |  | f2 | 1.350 | 0.260 | 0.066 |
|  | f3 | 0.109 | 0.745 | 0.006 |  | f3 | 0.144 | 0.709 | 0.008 |
|  | f4 | 0.056 | 0.816 | 0.003 |  | f4 | 0.069 | 0.796 | 0.004 |
|  | f5 | 0.176 | 0.679 | 0.009 |  | f5 | 0.189 | 0.669 | 0.010 |
|  | f6 | 0.680 | 0.420 | 0.035 |  | f6 | 0.724 | 0.405 | 0.037 |
|  | f7 | 0.950 | 0.342 | 0.048 |  | f7 | 0.994 | 0.331 | 0.050 |
|  | f8 | 0.685 | 0.418 | 0.035 |  | f8 | 0.673 | 0.422 | 0.034 |
| ipsiCont * anteriority | f1 | 0.345 | 0.564 | 0.018 | ipsiCont * anteriority | f1 | 0.469 | 0.502 | 0.024 |
|  | f2 | 3.362 | 0.082 | 0.150 |  | f2 | 3.932 | 0.062 | 0.171 |
|  | f3 | 6.342 | 0.021 | 0.250 |  | f3 | 6.733 | 0.018 | 0.262 |
|  | f4 | 4.485 | 0.048 | 0.191 |  | f4 | 4.344 | 0.051 | 0.186 |
|  | f5 | 3.130 | 0.093 | 0.141 |  | f5 | 3.088 | 0.095 | 0.140 |
|  | f6 | 2.720 | 0.116 | 0.125 |  | f6 | 2.779 | 0.112 | 0.128 |
|  | f7 | 1.988 | 0.175 | 0.095 |  | f7 | 1.949 | 0.179 | 0.093 |
|  | f8 | 0.077 | 0.785 | 0.004 |  | f8 | 0.070 | 0.794 | 0.004 |
| ipsiCont * anteriority * ageGroup | f1 | 0.340 | 0.567 | 0.018 | ipsiCont * anteriority * ageGroup | f1 | 0.380 | 0.545 | 0.020 |
|  | f2 | 1.025 | 0.324 | 0.051 |  | f2 | 1.070 | 0.314 | 0.053 |
|  | f3 | 0.325 | 0.575 | 0.017 |  | f3 | 0.342 | 0.565 | 0.018 |
|  | f4 | 0.174 | 0.681 | 0.009 |  | f4 | 0.164 | 0.690 | 0.009 |
|  | f5 | 1.117 | 0.304 | 0.056 |  | f5 | 1.126 | 0.302 | 0.056 |
|  | f6 | 1.543 | 0.229 | 0.075 |  | f6 | 1.608 | 0.220 | 0.078 |
|  | f7 | 2.691 | 0.117 | 0.124 |  | f7 | 2.702 | 0.117 | 0.125 |
|  | f8 | 0.902 | 0.354 | 0.045 |  | f8 | 0.907 | 0.353 | 0.046 |

**Table 3: Experiment 2 - table supplement 3:** Results of the repeated measures ANOVA with three witin-subjects factors for the frequency decomposed AAM. f1 to f8 refer to the subbands in descending order of frequency. ipsiCont: ipsilateral vs. contralateral channel (with respect to the attended side). ageGroup: older vs. younger adults. anteriority: stimulus presentation from the front or back speakers. The left columns show the results when only the corresponding channel was used to reject artifacts, while the right columns display the results when artifact rejection additionally considers the M.SCM and HEOG.

| Subbands AAM |  |  |  |  |  |  |  |  |  |
| --- | --- | --- | --- | --- | --- | --- | --- | --- | --- |
| Artifact rejection based on channel only |  |  |  |  | Artifact rejection based on channel, M.SCM and HEOG |  |  |  |  |
| Source | Measure | F | Sig. | Partial Eta Squared | Source | Measure | F | Sig. | Partial Eta Squared |
| ipsiCont | f1 | 0.900 | 0.355 | 0.045 | ipsiCont | f1 | 1.277 | 0.273 | 0.063 |
|  | f2 | 1.870 | 0.187 | 0.090 |  | f2 | 2.311 | 0.145 | 0.108 |
|  | f3 | 3.594 | 0.073 | 0.159 |  | f3 | 4.020 | 0.059 | 0.175 |
|  | f4 | 7.114 | 0.015 | 0.272 |  | f4 | 7.664 | 0.012 | 0.287 |
|  | f5 | 8.231 | 0.010 | 0.302 |  | f5 | 8.488 | 0.009 | 0.309 |
|  | f6 | 6.789 | 0.017 | 0.263 |  | f6 | 7.065 | 0.016 | 0.271 |
|  | f7 | 4.333 | 0.051 | 0.186 |  | f7 | 4.293 | 0.052 | 0.184 |
|  | f8 | 5.408 | 0.031 | 0.222 |  | f8 | 5.851 | 0.026 | 0.235 |
| ipsiCont * ageGroup | f1 | 0.001 | 0.971 | 0.000 | ipsiCont * ageGroup | f1 | 0.015 | 0.904 | 0.001 |
|  | f2 | 0.228 | 0.639 | 0.012 |  | f2 | 0.434 | 0.518 | 0.022 |
|  | f3 | 0.043 | 0.837 | 0.002 |  | f3 | 0.146 | 0.707 | 0.008 |
|  | f4 | 0.000 | 0.983 | 0.000 |  | f4 | 0.011 | 0.917 | 0.001 |
|  | f5 | 0.102 | 0.752 | 0.005 |  | f5 | 0.095 | 0.762 | 0.005 |
|  | f6 | 0.707 | 0.411 | 0.036 |  | f6 | 0.719 | 0.407 | 0.036 |
|  | f7 | 1.751 | 0.201 | 0.084 |  | f7 | 1.778 | 0.198 | 0.086 |
|  | f8 | 0.182 | 0.674 | 0.010 |  | f8 | 0.120 | 0.733 | 0.006 |
| anteriority | f1 | 3.259 | 0.087 | 0.146 | anteriority | f1 | 2.831 | 0.109 | 0.130 |
|  | f2 | 3.080 | 0.095 | 0.139 |  | f2 | 2.694 | 0.117 | 0.124 |
|  | f3 | 3.828 | 0.065 | 0.168 |  | f3 | 3.526 | 0.076 | 0.157 |
|  | f4 | 4.262 | 0.053 | 0.183 |  | f4 | 4.287 | 0.052 | 0.184 |
|  | f5 | 3.190 | 0.090 | 0.144 |  | f5 | 2.992 | 0.100 | 0.136 |
|  | f6 | 1.795 | 0.196 | 0.086 |  | f6 | 1.592 | 0.222 | 0.077 |
|  | f7 | 1.872 | 0.187 | 0.090 |  | f7 | 1.869 | 0.188 | 0.090 |
|  | f8 | 19.224 | 0.000 | 0.503 |  | f8 | 19.582 | 0.000 | 0.508 |
| anteriority * ageGroup | f1 | 0.112 | 0.742 | 0.006 | anteriority * ageGroup | f1 | 0.214 | 0.649 | 0.011 |
|  | f2 | 0.005 | 0.944 | 0.000 |  | f2 | 0.052 | 0.822 | 0.003 |
|  | f3 | 0.012 | 0.916 | 0.001 |  | f3 | 0.038 | 0.847 | 0.002 |
|  | f4 | 0.051 | 0.824 | 0.003 |  | f4 | 0.054 | 0.819 | 0.003 |
|  | f5 | 0.001 | 0.970 | 0.000 |  | f5 | 0.001 | 0.980 | 0.000 |
|  | f6 | 0.032 | 0.860 | 0.002 |  | f6 | 0.010 | 0.921 | 0.001 |
|  | f7 | 0.002 | 0.967 | 0.000 |  | f7 | 0.001 | 0.974 | 0.000 |
|  | f8 | 0.765 | 0.393 | 0.039 |  | f8 | 0.807 | 0.380 | 0.041 |
| ipsiCont * anteriority | f1 | 0.039 | 0.846 | 0.002 | ipsiCont * anteriority | f1 | 0.005 | 0.946 | 0.000 |
|  | f2 | 0.070 | 0.794 | 0.004 |  | f2 | 0.000 | 0.997 | 0.000 |
|  | f3 | 0.508 | 0.485 | 0.026 |  | f3 | 1.018 | 0.326 | 0.051 |
|  | f4 | 3.528 | 0.076 | 0.157 |  | f4 | 3.781 | 0.067 | 0.166 |
|  | f5 | 4.482 | 0.048 | 0.191 |  | f5 | 4.865 | 0.040 | 0.204 |
|  | f6 | 3.661 | 0.071 | 0.162 |  | f6 | 4.322 | 0.051 | 0.185 |
|  | f7 | 1.812 | 0.194 | 0.087 |  | f7 | 1.993 | 0.174 | 0.095 |
|  | f8 | 0.213 | 0.649 | 0.011 |  | f8 | 0.220 | 0.644 | 0.011 |
| ipsiCont * anteriority * ageGroup | f1 | 1.304 | 0.268 | 0.064 | ipsiCont * anteriority * ageGroup | f1 | 1.504 | 0.235 | 0.073 |
|  | f2 | 0.192 | 0.666 | 0.010 |  | f2 | 0.100 | 0.755 | 0.005 |
|  | f3 | 0.001 | 0.974 | 0.000 |  | f3 | 0.107 | 0.748 | 0.006 |
|  | f4 | 0.001 | 0.981 | 0.000 |  | f4 | 0.048 | 0.829 | 0.003 |
|  | f5 | 0.037 | 0.850 | 0.002 |  | f5 | 0.112 | 0.742 | 0.006 |
|  | f6 | 0.075 | 0.788 | 0.004 |  | f6 | 0.196 | 0.663 | 0.010 |
|  | f7 | 0.173 | 0.682 | 0.009 |  | f7 | 0.175 | 0.680 | 0.009 |
|  | f8 | 0.286 | 0.599 | 0.015 |  | f8 | 0.282 | 0.602 | 0.015 |

**Table 4: Experiment 2 - table supplement 4:** Results of the repeated measures ANOVA with three within-subjects factors for the frequency decomposed TAM. f1 to f8 refer to the subbands in descending order of frequency. ipsiCont: ipsilateral vs. contralateral channel (with respect to the attended side). ageGroup: older vs. younger adults. anteriority: stimulus presentation from the front or back speakers. The left columns show the results when only the corresponding channel was used to reject artifacts, while the right columns display the results when artifact rejection additionally considers the M.SCM and HEOG.

| Subbands TAM |  |  |  |  |  |  |  |  |  |
| --- | --- | --- | --- | --- | --- | --- | --- | --- | --- |
| Artifact rejection based on channel only |  |  |  |  | Artifact rejection based on channel, M.SCM and HEOG |  |  |  |  |
| Source | Mean Square | F | Sig. | Partial Eta Squared | Source | Measure | F | Sig. | Partial Eta Squared |
| ipsiCont | 3.444E-5 | 0.006 | 0.937 | 0.000 | ipsiCont | f1 | 0.012 | 0.915 | 0.001 |
|  | 000 | 0.005 | 0.946 | 0.000 |  | f2 | 0.009 | 0.927 | 0.000 |
|  | 003 | 0.064 | 0.803 | 0.003 |  | f3 | 0.054 | 0.818 | 0.003 |
|  | 017 | 0.308 | 0.585 | 0.016 |  | f4 | 0.277 | 0.605 | 0.014 |
|  | 024 | 0.461 | 0.505 | 0.024 |  | f5 | 0.466 | 0.503 | 0.024 |
|  | 025 | 0.558 | 0.464 | 0.029 |  | f6 | 0.571 | 0.459 | 0.029 |
|  | 016 | 0.381 | 0.544 | 0.020 |  | f7 | 0.390 | 0.540 | 0.020 |
|  | 010 | 1.558 | 0.227 | 0.076 |  | f8 | 1.434 | 0.246 | 0.070 |
| ipsiCont * ageGroup | 001 | 0.220 | 0.645 | 0.011 | ipsiCont * ageGroup | f1 | 0.270 | 0.609 | 0.014 |
|  | 003 | 0.087 | 0.771 | 0.005 |  | f2 | 0.093 | 0.763 | 0.005 |
|  | 000 | 0.004 | 0.948 | 0.000 |  | f3 | 0.008 | 0.930 | 0.000 |
|  | 7.160E-5 | 0.001 | 0.971 | 0.000 |  | f4 | 0.004 | 0.949 | 0.000 |
|  | 002 | 0.041 | 0.842 | 0.002 |  | f5 | 0.043 | 0.838 | 0.002 |
|  | 006 | 0.133 | 0.719 | 0.007 |  | f6 | 0.146 | 0.707 | 0.008 |
|  | 003 | 0.070 | 0.794 | 0.004 |  | f7 | 0.077 | 0.784 | 0.004 |
|  | 000 | 0.036 | 0.851 | 0.002 |  | f8 | 0.091 | 0.767 | 0.005 |
| anteriority | 044 | 6.104 | 0.023 | 0.243 | anteriority | f1 | 6.063 | 0.024 | 0.242 |
|  | 167 | 5.953 | 0.025 | 0.239 |  | f2 | 5.635 | 0.028 | 0.229 |
|  | 311 | 7.169 | 0.015 | 0.274 |  | f3 | 6.688 | 0.018 | 0.260 |
|  | 385 | 8.129 | 0.010 | 0.300 |  | f4 | 7.834 | 0.011 | 0.292 |
|  | 370 | 8.176 | 0.010 | 0.301 |  | f5 | 7.666 | 0.012 | 0.287 |
|  | 326 | 7.942 | 0.011 | 0.295 |  | f6 | 7.638 | 0.012 | 0.287 |
|  | 228 | 7.330 | 0.014 | 0.278 |  | f7 | 7.287 | 0.014 | 0.277 |
|  | 317 | 42.497 | 0.000 | 0.691 |  | f8 | 42.834 | 0.000 | 0.693 |
| anteriority * ageGroup | 003 | 0.377 | 0.546 | 0.019 | anteriority * ageGroup | f1 | 0.391 | 0.539 | 0.020 |
|  | 032 | 1.138 | 0.299 | 0.057 |  | f2 | 1.012 | 0.327 | 0.051 |
|  | 073 | 1.682 | 0.210 | 0.081 |  | f3 | 1.402 | 0.251 | 0.069 |
|  | 072 | 1.517 | 0.233 | 0.074 |  | f4 | 1.343 | 0.261 | 0.066 |
|  | 036 | 0.793 | 0.384 | 0.040 |  | f5 | 0.637 | 0.435 | 0.032 |
|  | 018 | 0.446 | 0.512 | 0.023 |  | f6 | 0.355 | 0.558 | 0.018 |
|  | 016 | 0.516 | 0.481 | 0.026 |  | f7 | 0.509 | 0.484 | 0.026 |
|  | 013 | 1.709 | 0.207 | 0.083 |  | f8 | 1.688 | 0.209 | 0.082 |
| ipsiCont * anteriority | 000 | 0.170 | 0.685 | 0.009 | ipsiCont * anteriority | f1 | 0.206 | 0.655 | 0.011 |
|  | 002 | 0.380 | 0.545 | 0.020 |  | f2 | 0.450 | 0.510 | 0.023 |
|  | 008 | 0.918 | 0.350 | 0.046 |  | f3 | 1.079 | 0.312 | 0.054 |
|  | 006 | 0.533 | 0.474 | 0.027 |  | f4 | 0.598 | 0.449 | 0.031 |
|  | 002 | 0.175 | 0.681 | 0.009 |  | f5 | 0.249 | 0.624 | 0.013 |
|  | 001 | 0.075 | 0.787 | 0.004 |  | f6 | 0.109 | 0.745 | 0.006 |
|  | 005 | 0.658 | 0.427 | 0.033 |  | f7 | 0.735 | 0.402 | 0.037 |
|  | 001 | 0.581 | 0.455 | 0.030 |  | f8 | 0.608 | 0.445 | 0.031 |
| ipsiCont * anteriority * ageGroup | 000 | 0.519 | 0.480 | 0.027 | ipsiCont * anteriority * ageGroup | f1 | 0.482 | 0.496 | 0.025 |
|  | 003 | 0.571 | 0.459 | 0.029 |  | f2 | 0.529 | 0.476 | 0.027 |
|  | 004 | 0.428 | 0.521 | 0.022 |  | f3 | 0.341 | 0.566 | 0.018 |
|  | 000 | 0.038 | 0.848 | 0.002 |  | f4 | 0.020 | 0.888 | 0.001 |
|  | 000 | 0.022 | 0.884 | 0.001 |  | f5 | 0.061 | 0.808 | 0.003 |
|  | 000 | 0.011 | 0.916 | 0.001 |  | f6 | 0.038 | 0.848 | 0.002 |
|  | 001 | 0.103 | 0.751 | 0.005 |  | f7 | 0.130 | 0.723 | 0.007 |
|  | 000 | 0.186 | 0.671 | 0.010 |  | f8 | 0.172 | 0.683 | 0.009 |

**Table 5: Experiment 2 - table supplement 5:** Results of the repeated measures ANOVA with three within-subjects factors for the frequency decomposed SAM. f1 to f8 refer to the subbands in descending order of frequency. ipsiCont: ipsilateral vs. contralateral channel (with respect to the attended side). ageGroup: older vs. younger adults. anteriority: stimulus presentation from the front or back speakers. The left columns show the results when only the corresponding channel was used to reject artifacts, while the right columns display the results when artifact rejection additionally considers the M.SCM and HEOG.

| Subbands SAM |  |  |  |  |  |  |  |  |  |
| --- | --- | --- | --- | --- | --- | --- | --- | --- | --- |
| Artifact rejection based on channel only |  |  |  |  | Artifact rejection based on channel, M.SCM and HEOG |  |  |  |  |
| Source | Measure | F | Sig. | Partial Eta Squared | Source | Measure | F | Sig. | Partial Eta Squared |
| ipsiCont | f1 | 10.582 | 0.004 | 0.358 | ipsiCont | f1 | 11.524 | 0.003 | 0.378 |
|  | f2 | 15.800 | 0.001 | 0.454 |  | f2 | 17.718 | 0.000 | 0.483 |
|  | f3 | 16.313 | 0.001 | 0.462 |  | f3 | 16.499 | 0.001 | 0.465 |
|  | f4 | 19.565 | 0.000 | 0.507 |  | f4 | 20.312 | 0.000 | 0.517 |
|  | f5 | 24.001 | 0.000 | 0.558 |  | f5 | 24.642 | 0.000 | 0.565 |
|  | f6 | 23.512 | 0.000 | 0.553 |  | f6 | 23.446 | 0.000 | 0.552 |
|  | f7 | 32.793 | 0.000 | 0.633 |  | f7 | 33.528 | 0.000 | 0.638 |
|  | f8 | 10.776 | 0.004 | 0.362 |  | f8 | 11.013 | 0.004 | 0.367 |
| ipsiCont * ageGroup | f1 | 0.005 | 0.944 | 0.000 | ipsiCont * ageGroup | f1 | 0.092 | 0.765 | 0.005 |
|  | f2 | 0.681 | 0.420 | 0.035 |  | f2 | 1.407 | 0.250 | 0.069 |
|  | f3 | 1.045 | 0.320 | 0.052 |  | f3 | 1.483 | 0.238 | 0.072 |
|  | f4 | 0.999 | 0.330 | 0.050 |  | f4 | 1.363 | 0.257 | 0.067 |
|  | f5 | 1.279 | 0.272 | 0.063 |  | f5 | 1.556 | 0.227 | 0.076 |
|  | f6 | 1.201 | 0.287 | 0.059 |  | f6 | 1.307 | 0.267 | 0.064 |
|  | f7 | 0.887 | 0.358 | 0.045 |  | f7 | 0.972 | 0.337 | 0.049 |
|  | f8 | 0.189 | 0.668 | 0.010 |  | f8 | 0.209 | 0.653 | 0.011 |
| anteriority | f1 | 5.928 | 0.025 | 0.238 | anteriority | f1 | 4.908 | 0.039 | 0.205 |
|  | f2 | 7.385 | 0.014 | 0.280 |  | f2 | 6.710 | 0.018 | 0.261 |
|  | f3 | 10.718 | 0.004 | 0.361 |  | f3 | 10.282 | 0.005 | 0.351 |
|  | f4 | 12.880 | 0.002 | 0.404 |  | f4 | 12.693 | 0.002 | 0.400 |
|  | f5 | 12.391 | 0.002 | 0.395 |  | f5 | 12.424 | 0.002 | 0.395 |
|  | f6 | 12.730 | 0.002 | 0.401 |  | f6 | 12.864 | 0.002 | 0.404 |
|  | f7 | 12.338 | 0.002 | 0.394 |  | f7 | 12.590 | 0.002 | 0.399 |
|  | f8 | 71.990 | 0.000 | 0.791 |  | f8 | 74.329 | 0.000 | 0.796 |
| anteriority * ageGroup | f1 | 0.049 | 0.828 | 0.003 | anteriority * ageGroup | f1 | 0.321 | 0.577 | 0.017 |
|  | f2 | 0.041 | 0.842 | 0.002 |  | f2 | 0.008 | 0.929 | 0.000 |
|  | f3 | 0.001 | 0.980 | 0.000 |  | f3 | 0.001 | 0.974 | 0.000 |
|  | f4 | 0.100 | 0.756 | 0.005 |  | f4 | 0.078 | 0.783 | 0.004 |
|  | f5 | 0.153 | 0.700 | 0.008 |  | f5 | 0.116 | 0.737 | 0.006 |
|  | f6 | 0.088 | 0.770 | 0.005 |  | f6 | 0.059 | 0.811 | 0.003 |
|  | f7 | 0.069 | 0.796 | 0.004 |  | f7 | 0.089 | 0.769 | 0.005 |
|  | f8 | 0.709 | 0.410 | 0.036 |  | f8 | 0.744 | 0.399 | 0.038 |
| ipsiCont * anteriority | f1 | 1.320 | 0.265 | 0.065 | ipsiCont * anteriority | f1 | 0.601 | 0.448 | 0.031 |
|  | f2 | 0.013 | 0.910 | 0.001 |  | f2 | 0.213 | 0.650 | 0.011 |
|  | f3 | 0.165 | 0.689 | 0.009 |  | f3 | 0.302 | 0.589 | 0.016 |
|  | f4 | 0.426 | 0.522 | 0.022 |  | f4 | 0.504 | 0.486 | 0.026 |
|  | f5 | 0.158 | 0.695 | 0.008 |  | f5 | 0.189 | 0.669 | 0.010 |
|  | f6 | 0.039 | 0.845 | 0.002 |  | f6 | 0.047 | 0.831 | 0.002 |
|  | f7 | 0.009 | 0.926 | 0.000 |  | f7 | 0.004 | 0.949 | 0.000 |
|  | f8 | 0.412 | 0.529 | 0.021 |  | f8 | 0.371 | 0.550 | 0.019 |
| ipsiCont * anteriority * ageGroup | f1 | 1.466 | 0.241 | 0.072 | ipsiCont * anteriority * ageGroup | f1 | 1.187 | 0.290 | 0.059 |
|  | f2 | 0.713 | 0.409 | 0.036 |  | f2 | 0.262 | 0.614 | 0.014 |
|  | f3 | 0.012 | 0.915 | 0.001 |  | f3 | 0.010 | 0.923 | 0.001 |
|  | f4 | 0.271 | 0.609 | 0.014 |  | f4 | 0.379 | 0.545 | 0.020 |
|  | f5 | 0.851 | 0.368 | 0.043 |  | f5 | 0.959 | 0.340 | 0.048 |
|  | f6 | 1.455 | 0.243 | 0.071 |  | f6 | 1.536 | 0.230 | 0.075 |
|  | f7 | 0.880 | 0.360 | 0.044 |  | f7 | 0.906 | 0.353 | 0.046 |
|  | f8 | 0.393 | 0.538 | 0.020 |  | f8 | 0.466 | 0.503 | 0.024 |

## Footnotes

1 A cognitive neuroscientist commented to the author S.A.H. that “Twice in my life, I have had the peculiar experience of nearly having my eyeglasses fall off because my right ear ‘twitched’ very strongly in response to an unexpected sound in the periphery” (personal communication, Theodore A. Bell, 13 October, 2015)

